# scMultiSim: simulation of multi-modality single cell data guided by cell-cell interactions and gene regulatory networks

**DOI:** 10.1101/2022.10.15.512320

**Authors:** Hechen Li, Ziqi Zhang, Michael Squires, Xi Chen, Xiuwei Zhang

## Abstract

Simulated single-cell data is essential for designing and evaluating computational methods in the absence of experimental ground truth. Existing simulators typically focus on modeling one or two specific biological factors or mechanisms that affect the output data, which limits their capacity to simulate the complexity and multi-modality in real data. Here, we present scMultiSim, an *in silico* simulator that generates multi-modal single-cell data, including gene expression, chromatin accessibility, RNA velocity, and spatial cell locations while accounting for the relationships between modalities. scMultiSim jointly models various biological factors that affect the output data, including cell identity, within-cell gene regulatory networks (GRNs), cell-cell interactions (CCIs), and chromatin accessibility, while also incorporating technical noises. Moreover, it allows users to adjust each factor’s effect easily. We validated scMultiSim’s simulated biological effects and demonstrated its applications by benchmarking a wide range of computational tasks, including cell clustering and trajectory inference, multi-modal and multi-batch data integration, RNA velocity estimation, GRN inference and CCI inference using spatially resolved gene expression data. Compared to existing simulators, scMultiSim can benchmark a much broader range of existing computational problems and even new potential tasks.

## Introduction

In recent years, technologies that profile the transcriptome and other modalities (multi-omics) of single cell have brought remarkable advances in our understanding of cellular mechanisms [61]. For example, technologies have enabled the joint profiling of chromatin accessibility and gene expression data [10; 8; 41], as well as the measurement of surface protein abundance alongside transcriptome [56; 47]. Additionally, spatial locations of cells can be measured together with transcriptome profiles using imaging-based [52; 19; 63] or sequencing-based [55; 50] technologies.

The advent of single-cell multi-omics data has facilitated a more comprehensive understanding of cellular states, and more importantly, allowed researchers to explore the relationships between modalities and the causality across hierarchies [18]. Prior to the availability of single cell multi-omics data, gene regulatory network (GRN) inference methods were developed using only single-cell RNA sequencing (scRNA-seq) data [48]. However, these methods mainly focused on transcription factors (TFs) as the sole factor affecting gene expressions. In reality, the observed gene-expression data is affected by multiple factors, such as the chromatin accessibility of corresponding regions. Consequently, newer methods utilizing both scRNA-seq and scATAC-seq data have been developed to infer GRNs [30; 62; 68]. Similarly, there has been a surge in the development of other computational tools that harness multi-modality information. For instance, Cell-Cell Interaction (CCI) inference methods seek to utilize both the gene expression and the spatial location modalities [16; 53; 5; 6] to learn the interactions with a lower false-positive rate than those using only scRNA-seq data [4; 26; 29]. Data integration methods combine multi-omics data to obtain a wholistic view of cells [58; 64; 1; 70; 36]. Moreover, RNA velocity can be inferred from unspliced and spliced counts using scRNA-seq data to indicate the near-future state of each cell [35; 3]. Recently, methods have also been proposed to infer RNA velocity from jointly profiled chromatin accessibility and transcriptomics data [38].

Overall, a large number of computational methods have been developed using scRNA-seq data or single cell multi- and spatial-omics data [66]. However, the scarcity of *ground truth* in experimental data makes it difficult to evaluate the performance of proposed computational methods. To address this, *de novo* simulators have been widely used to evaluate the accuracy of computational methods by generating data that models biological mechanisms and provides ground truth for benchmarking. SymSim [69], for example, provides ground truth cell identity and gene identity and thus can benchmark clustering, trajectory inference and differential expression detection. SERGIO [15], BEELINE [48] and dyngen [7] can simulate scRNA-seq data with given ground truth GRNs for testing GRN inference methods; while SERGIO, dyngen and VeloSim [71] can provide ground truth RNA velocity for testing RNA velocity inference methods. mistyR [60] generates single cell gene expression data from a given CCI network and can test CCI inference methods. With the *de novo* simulators, users can easily control the input parameters and obtain the exact ground truth. In addition to *de novo* simulators, Crowell *et al* [12] discussed another category of single cell data simulators, namely the reference-based methods, which learn a generative model from a given real dataset and generate synthetic data [13; 59; 54; 2]. By design, these methods can output datasets that mimic the input reference data, but their flexibility can be limited by the specific reference dataset. Although they can provide ground truth cluster labels using annotations in the reference dataset or pre-determined labels during the simulation, none of the reference-based methods provides ground truth that is rarely available via domain knowledge, like GRNs, CCIs, or RNA velocity.

We consider that a desirable single cell simulator should meet several criteria: (1) it should generate as many modalities as possible to best represent a cell; (2) it should model as many biological factors and mechanisms that affect the output data as possible so that the output data has realistic complexity; and (3) it should provide ground truth of the biological factors to benchmark various computational methods. Most existing simulators generate only scRNA-seq data, and some generate only scATAC-seq data [44; 37]. Among the few ones that can generate multiple modalities, dyngen and SERGIO output unspliced and spliced counts with ground truth RNA velocity, while a reference-based simulator scDesign3 [54] can generate two modalities each with high dimensionality (*eg*. scRNA-seq and DNA methylation data), or one high-dimensional modality (*eg*. scRNA-seq) and spatial location data depending on the input reference dataset (Table S1).

In terms of the biological factors modeled in the simulator, existing *de novo* simulators model only one or a small subset of the following biological factors that affect gene expression in a cell: cell identity (cluster labels or positions on cell trajectories), chromatin accessibility, GRNs, and CCIs (Table S1). Data generated by reference-based simulators can inherently have these effects but it is challenging to obtain the ground truth of the biological factors, thus unable to measure the accuracy of a computational method.

In this paper, we present scMultiSim, a unified framework that models *all* the above biological factors as well as technical variations including sequencing noise and batch effect (Fig. 1a). For each single cell, it outputs the following modalities: unspliced and spliced mRNA counts, chromatin accessibility, and spatial location, while considering the cross-modality relationships. “Chromatin accessibility” is both an output modality (also called the scATAC-seq modality) and a biological factor that affects other output data (it affects the gene expression modality). scMultiSim provides ground truth information on cell identity (in terms of cell populations), RNA velocity, GRNs and CCIs, as well as relationships between chromatin accessibility and transcriptome data. Therefore, with one dataset, it can be used to evaluate methods for various computational tasks including clustering or trajectory inference, multi-modal and multi-batch data integration, RNA velocity estimation, GRN inference and CCI inference. Moreover, scMultiSim allows the users to adjust the effect of each biological factor on the output data, enabling them to investigate how the methods’ performance is affected by each factor when evaluating methods for a specific task. We present a comparison between scMultiSim and existing multi-modal simulators in Table S1. To our knowledge, scMultiSim is the most versatile simulator to date in terms of its benchmarking applications.

**Figure 1.**
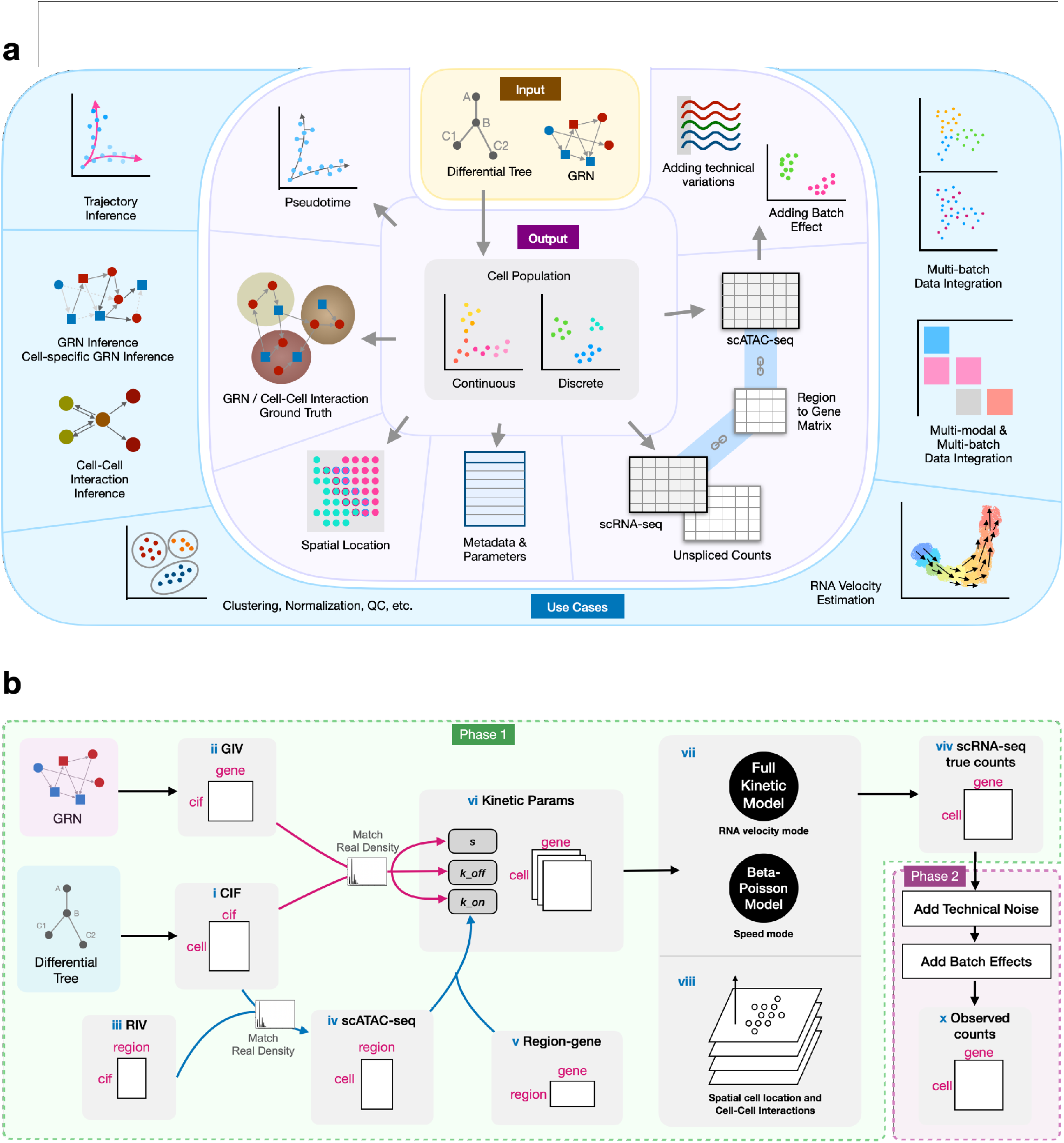
Overview of scMultiSim. **(a)**: The input, output, and use cases. The minimal required input is a cell differential tree describing the differentiation relationship of cell types. It controls the cell trajectory or clusters in the output. A user-input ground truth GRN is recommended to guide the simulation. Users can also provide ground truth for cell-cell interaction and control each simulated biological effects using various parameters. **(b)**: The overall structure of scMultiSim. The scATAC-seq data (iv) is firstly generated using CIF (i) and RIV (iii). The kinetic parameters used to generate scRNA-seq data (vi) is prepared using GIV (ii), CIF (i) and the scATAC-seq data with (**v**) a region-to-gene matrix. Using the parameters, either the full kinetic model (when RNA velocity is required), or the Beta-Poisson model (when running speed matters) will be used to generate the scRNA-seq data (vii). scMultiSim uses a multiple-step approach that considers both time and space when CCI is enabled (viii). With the simulated true counts (viv), technical noise and batch effects can be added to obtain the observed counts (x).

## Results

In the following sections, we will provide a brief overview of the core concepts and the simulation process of scMultiSim. We will then demonstrate its capability to simulate multiple biological factors simultaneously by validating the effects of each factor on the output data. Furthermore, we will showcase the applications of scMultiSim by using it to benchmark a wide variety of computational tools.

### scMultiSim overview

#### The kinetic model and control of intrinsic noise

In general, scMultiSim runs the simulation in two phases (Fig. 1b). In the first phase, scMultiSim employs the widely-accepted kinetic model [46] to generate the true gene expression levels in cells (“true counts”). In the second phase, scMultiSim introduces technical variations (library preparation noise, batch effects, etc) and generate scRNA-seq and scATAC-seq data that are statistically comparable to real data (“observed counts”). To model cellular heterogeneity and gene regulation effects, scMultiSim introduces two main concepts: *Cell Identity Factors* (CIFs) and *Gene Identity Vectors* (GIVs) (Fig. 1b (i, ii)). Biological factors, including cell population (cell identity), GRNs, and CCIs, are encoded in CIFs and GIVs (Fig. 2a). Additionally, to model single-cell chromatin accessibility, we also introduce Region Identity Vectors (RIVs, Fig. 1b(iii)). Further details on CIF, GIV and RIVs are provided in the next section.

**Figure 2.**
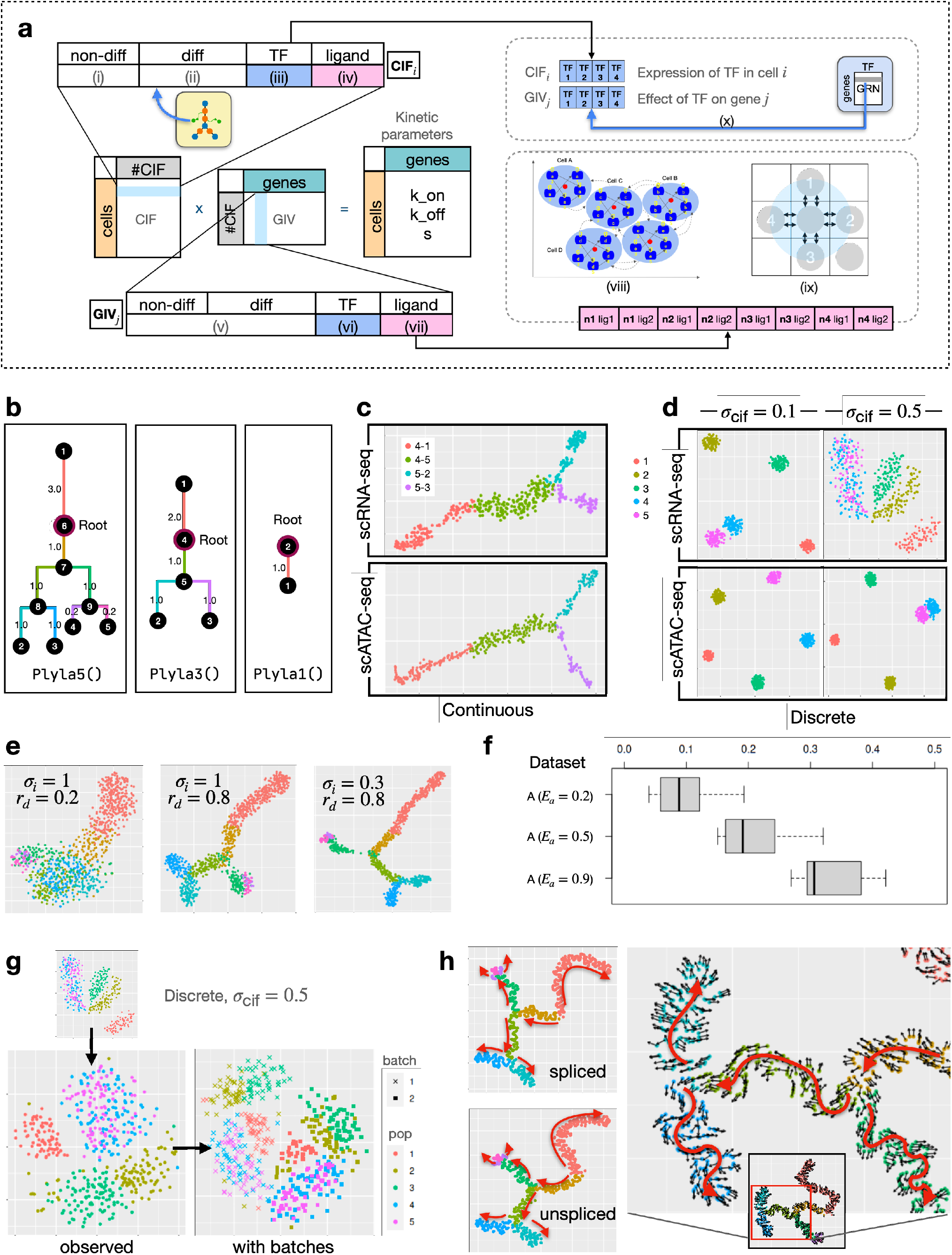
scMultiSim generates multi-modal single cell data from pre-defined cell clustering structure or trajectories. **(a)** The CIF and GIV matrix. We multiply the CIF and GIV matrix to get the cell*×*gene matrix for each kinetic parameter. CIFs and GIVs are divided into segments to encode different biological effects, where each segment encodes a certain type of biological factor. Cellular heterogeneity is modeled in the CIF, and regulation effects are encoded in the corresponding GIV vector. (viii) is the illustration of the cell-cell interactions and in-cell GRN in our model. (ix) is the grid system representing spatial locations of cells. A cell can have at most four neighbors (labeled 1-4) within a certain range (blue circle). The cell at the bottom right corner is not a neighbor of the center cell. (**b**) Three trees are provided by scMultiSim and used to produce the datasets. Phyla1 is a linear trejectory, while Phyla3 and Phyla5 has 3 and 5 leaves, respectively. (**c**) t-SNE visualization of the paired scRNA-seq and scATAC-seq data (without adding technical noise) from the main dataset MT3a (continuous populations following tree Phyla3), both having *n*_cell_ = *n*_gene_ = 500. (**d**) t-SNE visualization of the paired scRNA-seq and scATAC-seq data (without adding technical noise) from the main datasets MD3a and MD9a (discrete populations with five clusters, following tree Phyla5). (**e**) Additional results showing the effect of *σ*_*i*_ and *r*_*d*_ using datasets A. (**f**) Additional results exploring the ATAC effect parameter *E*_*a*_ using datasets A. Averaged Spearman correlation between scATAC-seq and scRNA-seq data for genes affected by one chromatin region, from 144 datasets using various parameters (*σ*_*i*_, *σ*_cif_, *r*_*d*_, continuous/discrete). All box plots in this article use the standard configuration, i.e., middle lines are medians, boxes represent the 1st and 3rd quartiles, and whiskers are *±*1.5 IQR. (**g**) The observed RNA counts in dataset MD9a with added technical noise and batch effects. (**h**) The spliced true counts, unspliced true counts, and the RNA velocity ground truth from dataset V. The velocity vectors point to the directions of differentiation indicated by red arrows, from the tree root to leaves.

When simulating single cell gene expression data, scMultiSim extends the idea of SymSim [69], where a kinetic model with three major parameters *k*_*on*_, *k*_*off*_, *s* was used to determine the expression pattern of a gene in a cell (Fig. 1b (vi)). In the kinetic model, a gene can switch between *on* and *off* states, with *k*_*on*_ and *k*_*off*_ be the rates of becoming *on* and *off*. When a gene is in the *on* state (which can be interpreted as promoter activation), mRNAs are synthesized at a rate *s* and degrade at a rate *d*. It is common to fix *d* at 1 and use the relative values for the other three parameters [43]. The kinetic parameters *k*_*on*_, *k*_*off*_, *s* are calculated from the CIF and GIV, as well as the corresponding scATAC-seq data (because chromatin accessibility is considered to affect gene expression). Since GIVs and CIFs encode information on cell identity, GRNs, and CCIs, the kinetic parameters thus capture the four biological factors that affect gene expression: cell identity, chromatin accessibility, GRNs, and CCIs.

The kinetic model used in scMultiSim provides two modes for generating true counts from the parameters, as shown in Fig. 1b (vii). The first mode is the full kinetic model, where genes undergo several cell cycles over time with *on*/*off* state changes, and the spliced/unspliced RNA counts are calculated. This mode provides the ground truth for RNA velocity since the RNA synthesis rate is known. The second mode is the Beta-Poisson model, which is equivalent to the kinetic model’s master equation [31], and is faster to run than the full kinetic model. The Beta-Poisson model is recommended when RNA velocity is not needed. In the Beta-Poisson model, scMultiSim also introduces an intrinsic noise parameter *σ*_*i*_ that controls the amount of intrinsic noise caused by the transcriptional burst and the snapshot nature of scRNA-seq data. This parameter allows users to examine the influence of intrinsic noise on the performance of the computational methods. The two modes and the *σ*_*i*_ parameter are described in Methods.

#### Modeling cellular heterogeneity and various biological effects

The design of *Cell Identity Factors (CIFs)* and *Gene Identity Vectors (GIVs)* allows scMultiSim to encode cell identities and gene-level mechanisms (such as GRNs and CCIs) into the kinetic parameters and thereby impact the gene expression levels. This design also provides easy ways to adjust the effect of each factor on the output gene expression data.

The CIF of a cell is a 1D vector representing various biological factors that contributes to cellular heterogeneity, such as the cell condition (*e*.*g*. treated or untreated), or the expression of key TFs. The GIV of a gene act as the weights of the corresponding factors in the CIF, representing how strongly the corresponding CIF affect the gene’s expression (Fig. 2a, Methods). By multiplying the CIF and GIV matrices, scMultiSim therefore generates a *n*_cell_ *× n*_gene_ matrix, which is the desired kinetic parameter matrix with the cell and gene factors encoded.

Each CIF vector and GIV vector consists of four segments, each representing one type of extrinsic variation. They encode biological factors including cell identity (cell population, *i*.*e*., the underlying cell trajectories or clusters), GRNs, and CCIs (Figs. 2a, S1a-b). We introduce the four segments in the following.

i. Non-differential CIFs (**non-diff-CIF**) model the inherent cellular heterogeneity. They represent various environmental factors or conditions that are shared across all cells and are sampled from a Gaussian distribution with standard deviation *σ*_cif_.
ii. Differential CIFs (**diff-CIF**) control the user-desired cell population. These are the biological conditions that are unique to certain cell types. These factors lead to different cell types in the data. For a heterogeneous cell population, cells have different development statuses and types. Values for diff-CIFs are used to represent these cell differential factors, which are generated based on the user-input cell differential tree. When generating data for cells from more than one cell type, the minimal user input of scMultiSim is the cell differential tree, which controls the cell types (for discrete populations) or trajectories (for continuous populations) in the output. The process of generating diff-CIFs is described in Methods.
iii. CIFs corresponding to Transcription Factors (**tf-CIF**) control the effects of GRNs. This segment, together with the TF segment in the GIV, model how a TF can affect expression of genes in the cell (Methods). Its length equals to the number of TFs. In other words, the GRN is encoded in the tf-CIFs and GIVs.
iv. CIFs corresponding to ligands from neighboring cells (**lig-CIF**) control the effect of CCI. If CCI simulation is enabled, this segment together with the ligand segment in the GIV of the receptor gene encodes the ground truth CCI between two cells. This encoding ensures that a ligand and its interacting receptor have correlated gene expression. A receptor can also interact with ligands of multiple neighbors (Fig. 2a (viii)). The GIV matrices are generated carefully considering the nature of the kinetic model (Methods).

#### The simulation process

Fig. 1b shows an overview of the simulation process. The scATAC-seq data is generated at first (Fig. 2b(iv)), because we consider that the chromatin accessibility of a cell affects its gene expression. The scATAC-seq data also follows a pre-defined clustering or trajectory structure represented by the input cell differentiation tree. Similar to the gene expression, we multiply the CIF with a Region Identity Vector (RIV) matrix, which represents the effect of each CIF on the accessibility of chromatin regions. Details on generating the scATAC-seq data are included in Methods. The scATAC-seq data affects scRNA-seq data through the *k*_*on*_ parameter, because chromatin accessibility controls the activated status of genes (Methods).

After obtaining all the kinetic parameters, scRNA-seq data can be generated in different modes: with or without CCIs and spatial locations, and with or without outputting RNA velocity data (Fig. 1b (vii, viii)). If the user specify to generate RNA velocity, the full kinetic model is used, where cells undergo several cycles before the spliced and unspliced counts are outputted (Methods). Otherwise, if the Beta-Poisson model is used, and the true counts are sampled from the Beta-Poisson distribution. In this mode, RNA velocity and unspliced count data are not outputted.

#### Simulating cell-cell interaction

If specified to generate spatial-aware single cell gene expression data including cell spatial locations and CCI effects, scMultiSim uses a multiple-step approach that considers both time and space (Fig. 1b (viii), Fig. S1c). The simulation consists of a series of steps, with each step representing a time point. Cells are placed in a grid (Fig. 2a (ix), Fig. S1d), and one cell is added to the grid at each step, representing a newborn cell. Users can use the parameter *p*_*n*_ to control the probability for the newborn cell to locate with cells of the same type (Methods). As experimental data cannot measure cells at previous time points, scMultiSim outputs data only for cells at the final time point, which contains the accumulated CCI effects during the cells’ developmental process.

To simulate CCI, scMultiSim requires a user-inputted list of ligand-receptor gene pairs that can potentially interact, which is called a ligand-receptor database. Users can input cell-type-level or single cell level CCI ground truth. If users do not provide ground truth CCIs, scMultiSim can randomly generate the ground truth from the ligand-receptor database.

#### Technical variations and batch effects

The steps described above belong to the first phase, which generates the “true” mRNA counts (and unspliced counts if RNA velocity mode is enabled) in the cells. In the second phase, scMultiSim simulates key experimental steps in wet labs that lead to technical noises in the data and output the observed scRNA-seq data. Batch effects can also be added to simulate datasets from a user-specified number of batches. Users can also control the amount of technical noise and batch effects between batches. These procedures are described in Methods. Next, we show the various output of scMultiSim and validate the effects present in the simulated data.

### Design of simulation and datasets

We have generated a comprehensive set of datasets using scMultiSim to demonstrate the effects of different parameter configurations and to benchmark computational methods. These datasets contain both *main* and *auxiliary* datasets. The main datasets consists of 144 datasets with varying configurations of important parameters, including *σ*_cif_ ∈ {0.1, 0.5*}, n*_cell_ ∈ {500, 800*}, n*_gene_ ∈ {110, 200, 500}, and three different cell trajectories. The *σ*_cif_ parameter controls the standard deviation of the CIF and affects the within-cluster or within-neighborhood heterogeneity between cells. These main datasets contain all effects scMultiSim can simulate: GRN, chromatin accessibility, cell-cell interaction, technical noise and batch effect. Thus, the 144 main datasets cover a wide range of variety, including different numbers of cells, genes, and trajectory shapes, to minimize potential bias and provide a more comprehensive benchmark of the computational methods.

As presented in Table 1, we label the main datasets with the following format: `M{p}{c}{s}`. The first letter M denotes the main dataset, followed by a letter `p` ∈ *{L, T, D}* that specifies the cell population as linear trajectory, tree trajectory or discrete, respectively. The number `c` ∈ [1, 12] denotes a particular configuration of *σ*_cif_, *n*_cell_, and *n*_gene_, while the last lowercase letter `s` ∈ *{a, b, c, d}* represents random seed 1-4. For instance, the dataset `MD5c` has a discrete cell population, *σ*_cif_ = 0.1, 800 cells, 200 genes and random seed 3.

**Table 1.**
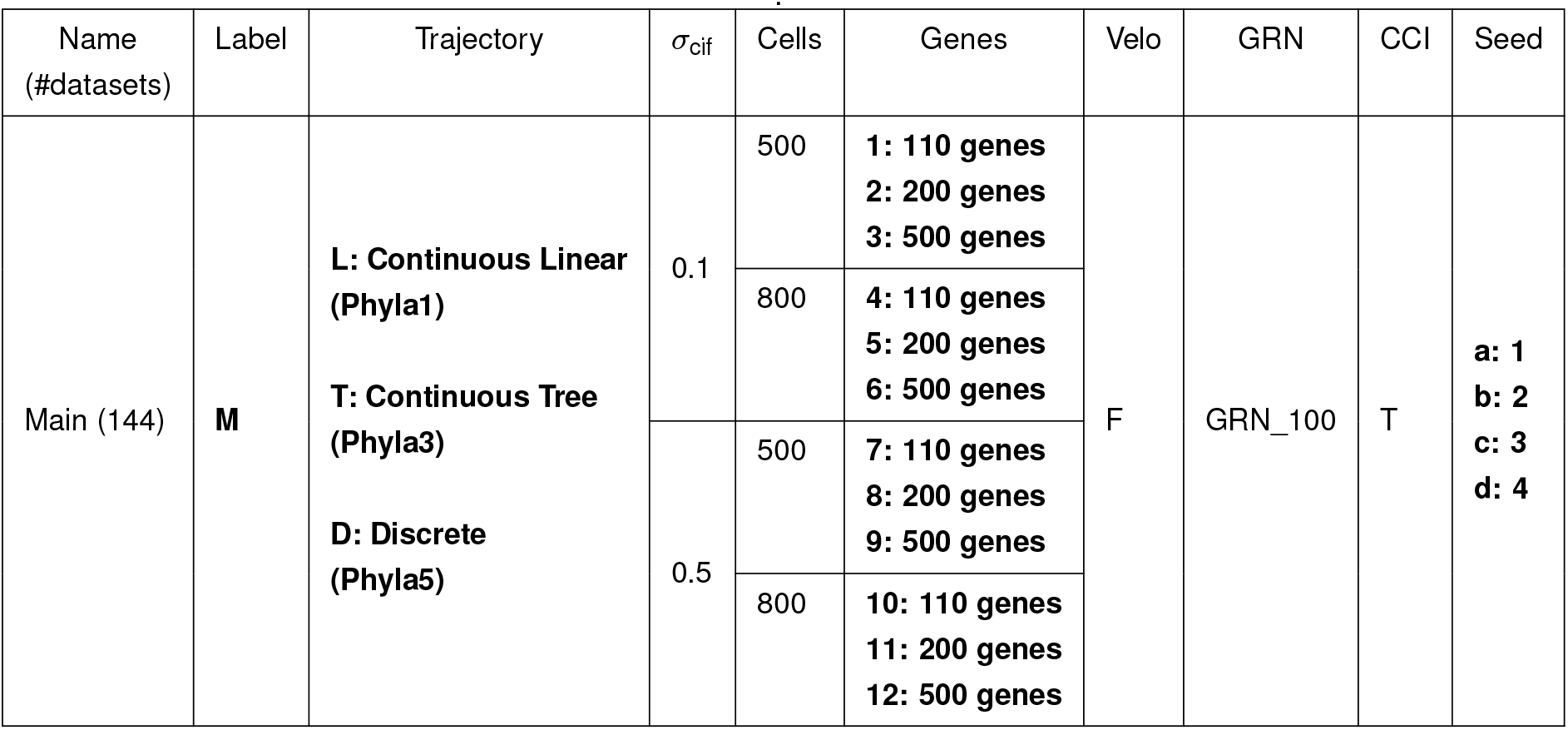
The main dataset contains 144 datasets with varying trajectory, *σ*_cif_,number of cells and genes. For each parameter configuration, four datasets are generated using different random seeds. We number the datasets for easy referencing in the text: starting with the letter M,then a letter {L,T,D} specifying the trajectory; followed by a number 1-12 identifying the configuration of *σ*_cif_, number of cells and genes; and last, a lowercase letter a-d indicating the random seed. For example, MD5c uses a discrete cell population, *σ*_cif_ = 0.1, 800 cells, 200 genes and random seed 3. Phyla1, Phyla3 and Phyla5 are the input tree structure used to generate the cell populations, and they are shown in Fig. 2b.

| Name<br>(#datasets) | Label | Trajectory | $\sigma_{\text{cif}}$ | Cells | Genes | Velo | GRN | CCI | Seed |
| --- | --- | --- | --- | --- | --- | --- | --- | --- | --- |
| Main (144) | **M** | **L: Continuous Linear**<br>**(Phyla1)**<br>**T: Continuous Tree**<br>**(Phyla3)**<br>**D: Discrete**<br>**(Phyla5)** | 0.1 | 500 | **1: 110 genes**<br>**2: 200 genes**<br>**3: 500 genes** | F | GRN_100 | T | **a: 1**<br>**b: 2**<br>**c: 3**<br>**d: 4** |
|  |  |  |  | 800 | **4: 110 genes**<br>**5: 200 genes**<br>**6: 500 genes** |  |  |  |  |
|  |  |  | 0.5 | 500 | **7: 110 genes**<br>**8: 200 genes**<br>**9: 500 genes** |  |  |  |  |
|  |  |  |  | 800 | **10: 110 genes**<br>**11: 200 genes**<br>**12: 500 genes** |  |  |  |  |

We have also generated auxiliary datasets with fewer types of effects and presented them in Table 2. These datasets allow us to explore the effect of other parameters and are compatible with computational methods that impose additional constraints on the input. In the remaining, we will primarily use the main datasets `M` for benchmarking and demonstration, while the auxiliary datasets will serve as additional and supplementary results.

**Table 2.** The auxiliary dataset and other datasets used in supplimental information.

| Name<br>(#datasets) | Label | Trajectory | $\sigma_{\text{cif}}$ | Cells | Genes | Velo | GRN | CCI | Seed | Other<br>Params |
| --- | --- | --- | --- | --- | --- | --- | --- | --- | --- | --- |
| Auxiliary (144) | **A** | Tree<br>(Phyla5) | 0.1, 0.5, 1 | 200 | 1000 | F | GRN_100 | F | 1,2 | $E_{\text{atac}} = 0.2, 0.5, 0.9$<br>$\sigma_i = 0.3, 1$<br>$r_d = 0.2, 0.8$ |
|  |  | Discrete<br>(5 clusters) |  |  |  |  |  |  |  |  |
| Velocity (72) | **V** | Tree<br>(Phyla5) | 0.1 | 500,750,1000 | 100,200,500 | T | GRN_100<br>N/A | T | 5-8 |  |
| Realistic (1) | **R** | Discrete<br>(10 clusters) | 0.1 | 523 | 200 | F | Inferred | F | 1 |  |
| Add. GRN (8) | **G** | Linear | 0.1 | 110 | 1000 | F | GRN_100 | F | 1-8 |  |
| Add. CCI (8) | **C** | Linear | 0.1 | 200 | 500 | F | Fig. S6 | T | 1-8 |  |

### scMultiSim generates multi-batch and multi-modality data from pre-defined clusters or trajectories

scMultiSim offers a key advantage in its ability to generate coupled scRNA-seq and scATAC-seq data while allowing users to control the shape of trajectories or clusters. It is accomplished by offering various parameters to control the structure of cell populations. First, the user can choose to generate “continuous” or “discrete” populations, and input a tree that represents the cell trajectories (in the case of “continuous” populations) or relationship between clusters (in the case of “discrete” populations). We name the tree “differentiation tree”. scMultiSim provides three example differentiation trees: Phyla1, Phyla3, and Phyla5, each having 1, 3, and 5 leaves, as illustrated in Fig. 2b. The main datasets were simulated using these trees (Table 1). From a differentiation tree, scMultiSim is able to generate both discrete and continuous cell populations (Fig. 2c). Then, users can use these three parameters: intrinsic noise *σ*_*i*_, CIF sigma *σ*_cif_ and Diff-to-nonDiff CIF ratio *r*_*d*_, to control how clean or noisy the population structure is in the data (Fig. 2c-e).

For the continuous population, we visualize a dataset `MT3a` generated using tree Phyla3 in Fig. 2c. We can observe that the trajectories corresponding to the input differentiation tree are clearly visible for both the scRNA-seq and the scATAC-seq modality. For the discrete population, we visualize dataset `MD3a` and `MD9a` generated with tree Phyla5 in Fig. 2d. The parameter *σ*_cif_ controls the standard deviation of the CIF, therefore with a smaller *σ*_cif_, the clusters are tighter and better separated from each other. We then used the auxiliary dataset `A` (Table 2) to explore the effect of the intrinsic noise parameter *σ*_*i*_ and *r*_*d*_, the ratio of number of diff-CIF to non-diff-CIFs. In Fig. 2e, we visualize the scRNA-seq modality generated using Phyla5 continuous mode with the same *σ*_cif_. With a smaller Diff-to-nonDiff CIF ratio *r*_*d*_, the trajectory is vague and more randomness is introduced, as the tree structure is encoded in the differential CIFs. With a smaller intrinsic noise *σ*_*i*_, a fraction of the expression value is directly calculated from kinetic parameters without sampling from the Poisson model;

As a result, the trajectory is more prominent. These patterns are much cleaner than real data because real data always has technical noise. We will show more results with technical noise in later sections and in Fig. S2.

#### Coupling between scATAC-seq and scRNA-seq data

In paired scATAC-seq and scRNA-seq data, these two data modalities are not independent of each other, as it is commonly considered that a gene’s expression level is affected by the chromatin accessibility of the corresponding regions. If a gene’s associated regions are accessible, this gene is more likely to be expressed. This mechanism can be naturally modeled in scMultiSim through the kinetic parameter *k*_*on*_ (Methods).

We provide a user-adjustable parameter, the ATAC-effect *E*_*a*_, to control the extent of scATAC-seq data’s effect on *k*_*on*_ (ranging from 0 and 1). In order to validate the connection between the scATAC-seq and scRNA-seq data, we calculate the mean Spearman correlation between these two modalities for genes that are controlled by one region in the scATAC-seq data. In Fig. 2f, we present the correlations under different *E*_*a*_ values. An averaged 0.2-0.3 correlation can be observed using the default value (0.5), and the correlation increases with higher values of *E*_*a*_. These results demonstrate that scMultiSim successfully models the connection between scATAC-seq and scRNA-seq data, enabling the generation of more realistic multi-omics datasets.

#### scMultiSim simulates technical noise and batch effect

The single cell gene expression data shown in Figs. 2c-f are “true” mRNA counts which do not have technical noise. scMultiSim can add technical noise including batch effects to the true counts to obtain observed counts (Methods). The amount of technical noise and batch effects can be adjusted through parameters, for example, the parameter *E*_batch_ can be used to control the amount of batch effect. Users can also specify the number of batches.

Fig. 2g shows the observed mRNA counts of dataset `MD9a` (true counts shown in Fig. 2d). The left plot shows data with one batch, and the right plot shows two batches. Technical noise and batch effects are also added to the scATAC-seq matrix. We further use the auxiliary dataset `A` to demonstrate the ability of scMultiSim to adjust the amount of technical noise and batch effect in both scRNA-seq and scATAC-seq modalities, in both continuous and discrete populations (Fig. S2). Here, we vary a main parameter for technical noise, *–*, which denotes the capture efficiency that affects the detection ability of the dataset. Lower *–* values correspond to poorer data quality.

### scMultiSim generates spliced and unspliced mRNA counts with ground truth RNA velocity

If RNA velocity simulation is enabled, the kinetic model outputs the velocity ground truth using the RNA splicing and degradation rates. The Phyla5 tree in Fig. 2b is used to generate the results in Fig. 2h. The figure shows both the true spliced and unspliced counts, as well as the ground truth RNA velocity averaged by *k* nearest neighbor (*k*NN), which can be used to benchmark RNA velocity estimation methods. The RNA velocity vectors follow the cell trajectory (backbone and directions shown in red), which is specified by the user-inputted differentiation tree.

### scMultiSim generates single cell gene expression data driven by GRNs and cell-cell interactions

The strength of scMultiSim also resides in its ability to incorporate the effect of GRN and CCI while preserving the pre-defined trajectory structures. In this section, we show that the GRN and CCI effects both exist in the simulated expression data. The main datasets (Table 1) used the 100-gene GRN from [15] as the ground truth GRN, which is visualized in Fig. 3a. We also incorporate CCIs by adding cross-cell ligand-receptor pairs to the within-cell GRNs. Specifically, we connect each cell’s gene 99,101-104 to a neighbor cell’s gene 91, 2, 6, 10 (TFs), and 8 (non-TF) in the GRN (green edges in Fig. 3a). Next, we use one dataset (`MT3a` with a tree trajectory, 500 genes, 500 cells, and *σ*_cif_ = 0.1) to inspect the simulated effects in detail (Fig. 3b-e).

**Figure 3.**
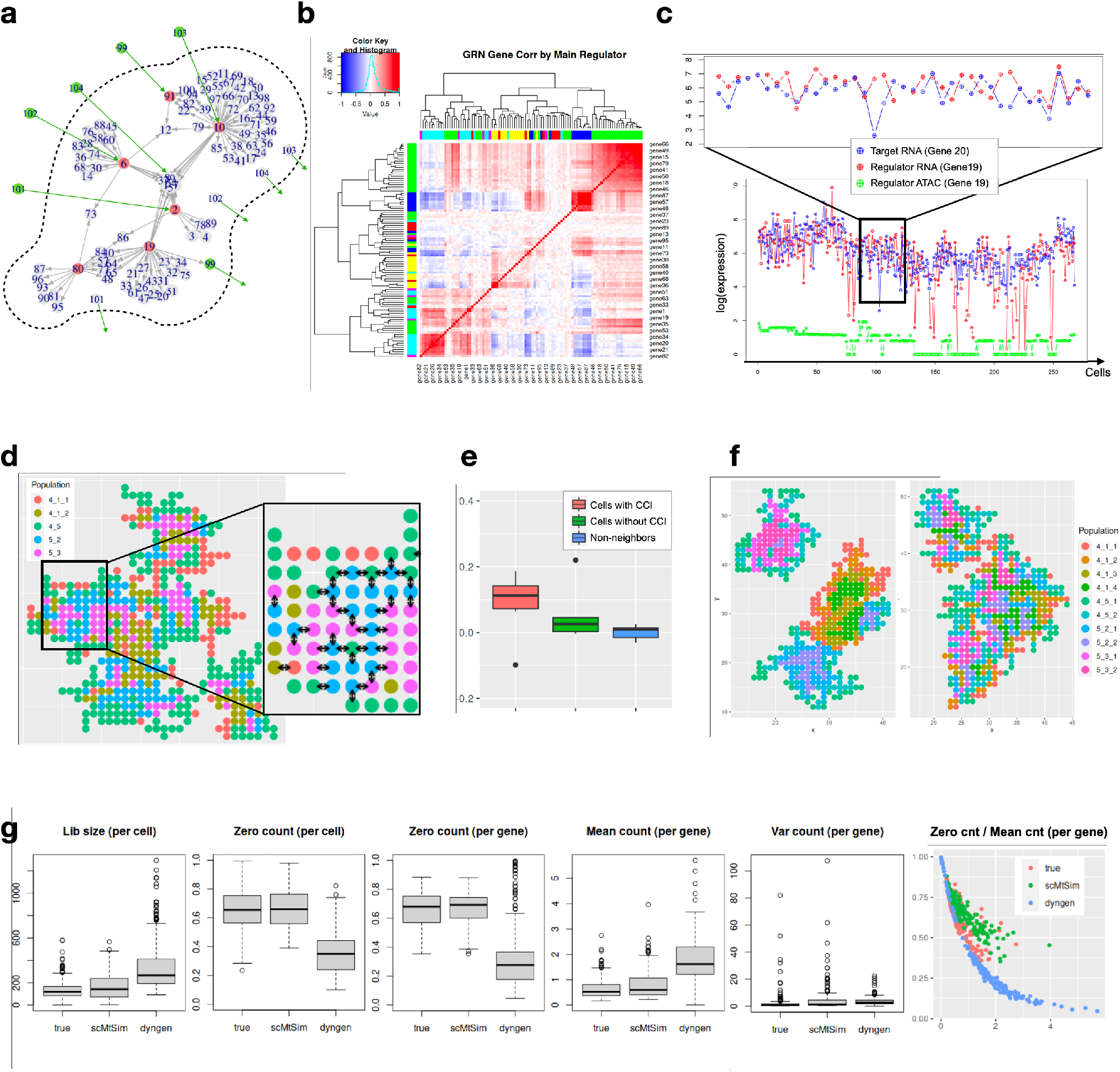
scMultiSim generates realistic single cell gene expression data driven by GRN and cell-cell interaction. (**a**) The GRN and CCIs used to generate the main datasets. Red nodes are TF genes and green nodes are ligand genes. Green edges are the added ligand-receptor pairs when simulating cell-cell interactions. (**b-e**) Results from dataset MT3a, which uses Phyla3, 500 genes, 500 cells and *σ*_cif_ = 0.1. (**b**) The gene module correlation heatmap. The color at left or top represents the regulating TF of the gene. Genes regulated by the same TF have higher correlations and tend to be grouped together. (**c**) The log-transformed expression of a specific TF-target gene pair (gene19-gene20) for all cells on one lineage (4-5-3 in Phyla3). Correlation between the TF and target expressions can be observed. We also show the chromatin accessibility level for the TF gene 19, averaged from the two corresponding chromatin regions of the gene. Significant lower expression of gene 19 can be observed when the chromatin is closed. (**d**) The spatial location of cells, where each color represents a cell type. Arrows between two cells indicates that CCI exists between them for a specific ligand-receptor pair (gene101-gene2). By default, most cell-cell interactions occur between different cell types. (**e**) Gene expression correlation between (1) neighboring cells with CCI, (2) neighboring cells with CCI, and (3) non-neighbor cells. Cells with CCI have higher correlations. (**f**) scMultiSim provides options to control the the cell layout. We show the results of 1200 cells using same-type probability *p*_*n*_ = 1.0 and 0.8, respectively. When *p*_*n*_ = 1.0, same-type cells tend to cluster together, while *p*_*n*_ = 0.8 introduces more randomness. (**g**) Comparison between a real dataset and simulated data using multiple statistical measurements (Methods). Parameters were adjusted to match the real distribution as close as possible.

#### GRN guided expression data

We illustrate the gene regulation effects for dataset `MT3a` using a gene module correlation heatmap as shown in Fig.3b. The clustered heatmap is constructed by computing pairwise Spearman correlations between the expression levels of all genes. Each color on the top or left side of the heatmap represents a TF in the GRN. The figure shows that gene modules regulated by the same TF (genes with the same color) tend to be clustered together and have higher correlations with each other. These results suggest the presence of GRN effects in the expression data. To further illustrate the regulatory effects, we plot the expression of a specific regulator-target pair (gene 19-20) along one lineage (4-5-3 in Phyla3) in Fig. 3c. The plot clearly shows a correlation between the expression levels of the regulator and target genes. Moreover, we plot the accessibility levels for the corresponding chromatin region of gene 19 in Fig.3c. The plot indicates that significant drops in gene 19’s expression occur when the related chromatin region is closed, providing further evidence for the regulatory effects of chromatin accessibility.

#### Cell spatial locations

scMultiSim provides convenient helper methods to visualize the cell spatial locations, as shown in Fig. 3d (dataset `MT3a`). For each ligand-receptor pair, arrows can be displayed between cells to show the direction of cell-cell interactions. We consider various biological scenarios when assigning the spatial location to each cell (Methods), for example, a newborn cell has a probability *p*_*n*_ of staying with a cell of the same type. Changing *p*_*n*_ allows us to generate different tissue layouts. In real data, how likely cells from the same cell type locate together depends on the tissue type, and scMultiSim provides *p*_*n*_ to tune this pattern. Fig. 3f shows the effect of varying *p*_*n*_. The left figure in the panel was generated with *p*_*n*_ = 1, showing strong spatial clustering of cells from the same cell type. The right figure in the panel was generated with *p*_*n*_ = 0.8, where cells from the same cell type are more spread out to enable more interactions across cell types.

#### Correlations between interacting ligands and receptors

scMultiSim simulates CCIs between single cells as well as between cell types. We validate the simulated CCI effects by comparing the correlations of expression levels between (i) neighboring cells with CCI, (ii) neighboring cells without CCI, and (iii) non-neighbor cells (Methods). As shown in Fig. 3e (using dataset `MT3a`), cells with CCI have an average pairwise correlation of 0.1, whereas cells without CCI exhibit approximately zero correlation, which is expected. We noticed that neighboring cells without CCI still have a slightly higher correlation compared to non-neighbor cells, which may be attributed to the dynamic nature of cell differentiation, where cells are evolving into new cell types over time, and CCI effects involved in an earlier cell type may remain in the final step.

### scMultiSim simulated datasets match real data

We show that scMultiSim’s output single cell gene expression data can statistically resemble real data. We used a spatially resolved single cell gene expression dataset measured with seqFISH+ technology [19; 16], and generated simulated data to match this real dataset (Methods). We used dyngen [7] as a baseline simulator to compare with, as it is also a *de novo* multi-modality simulator that shares a few functionalities with scMultiSim (Table S1). We compare the simulated data with real data in terms of the following properties: library size, zero counts per cell, zero counts per gene, mean count per gene, variation per gene, and the ratio between zero count and mean count per gene (Fig. 3g).

Fig. 3g shows that the library size, zero counts per cell, zero counts per gene and mean counts per gene simulated by scMultiSim are closer to that of real data than the dyngen simulated data, and both scMultiSim and dyngen are able to simulate data with realistic variation per gene. There is also usually a negative correlation between zero counts and mean counts in real data, and scMultiSim is able to simulate this relationship, matching well with the reference data.

### Benchmarking computational methods using scMultiSim

We next show that scMultiSim can be used to benchmark a board range of computational tasks in single cell genomics, including clustering, trajectory inference, multi-modal data integration, RNA velocity estimation, GRN inference and CCI inference using spatially resolved single cell gene expression data. Using scMultiSim, we studied the performance of several recent methods on each task, and also investigated the effect of particular parameters for some of the benchmarks. As far as we know, scMultiSim is the only simulator that can benchmark all these tasks. It is noteworthy that our intention is not to perform a comprehensive benchmarking analysis, but rather to show evidence of scMultiSim’s broad applications. We anticipate that these benchmarks can encourage forthcoming researchers to discover more use cases of scMultiSim.

### Benchmarking clustering and trajectory inference methods

We first applied scMultiSim to test methods for two classic problems: cell clustering and trajectory inference, using the scRNA-seq modality in our discrete main datasets (`MD`, Table 1). We tested five clustering methods, PCA-KMeans, CIDR [39], SC3 [33], TSCAN [28], and Seurat [23] (Fig. 4a). For each method and each dataset in the main datasets, we vary the parameter “number of clusters”. Since Seurat does not provide direct control over the number of clusters, we varied the resolution parameter instead and plotted using the number of clusters in the results. From Fig. 4a, all methods have the best performance when the cluster number is the true value. In general, Seurat and SC3 have better performance than the others, which is consistent with previous benchmarking [17]. TSCAN performs better than PCA-KMeans in our results which is not the case in [17]. We also show the comparison separately for *σ*_cif_ = 0.1 and *σ*_cif_ = 0.5 in Fig. S3a-b. Comparing Fig. S3a with Fig. S3b, the methods generally have higher ARI with a smaller *σ*_cif_, which is expected. Additionally, Seurat’s recommended resolution range (0.4-1.2) provides an accurate estimation of the number of clusters (Fig. S3c).

**Figure 4.**
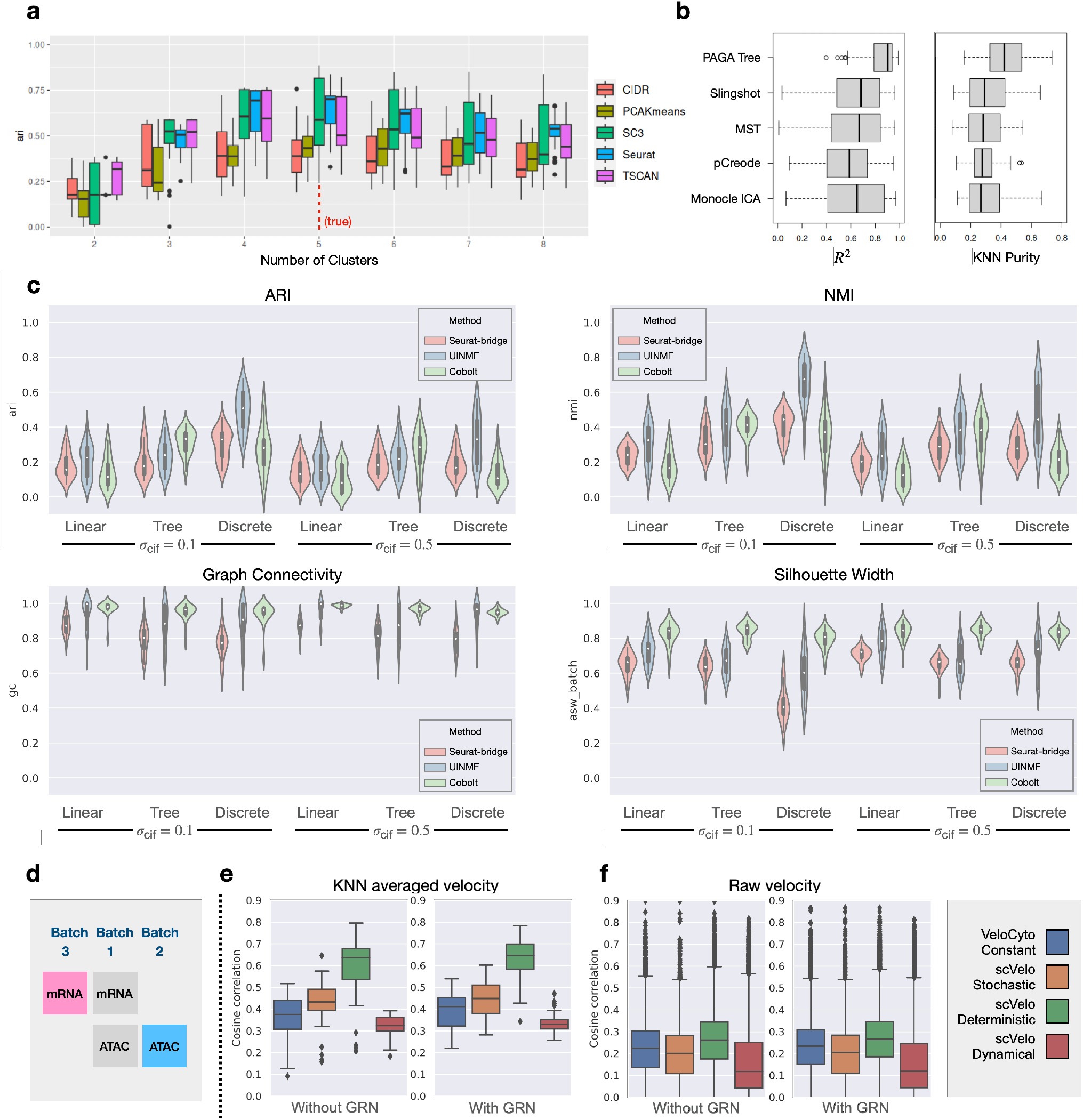
Benchmarking clustering, trajectory inference, multi-modal data integration and RNA velocity estimation methods. (**a**) Benchmarking clustering methods on dataset MD (discrete). Methods are grouped by number of clusters in the result. The vertical red dashed line shows the true number of clusters. A higher ARI indicates better clustering. (**b**) Benchmarking trajectory inference methods on dataset MT (continuous tree). Methods are evaluated based on their mean *R*^2^ and *k*NN purity on each lineage (higher is better). (**c**) Benchmarking multi-modal data integration methods. Metrics for the methods: ARI, NMI (higher = better at preserving cell identities), graph connectivity and average silhouette width of batch (higher = better merging batches). (**d**) The task illustration of multi-modal data integration. Only cells in batch 1 and 3 (pink and blue matrices) are used for evaluation. (**e**,**f**) Benchmarking RNA velocity estimation methods on auxiliary dataset V. The result is measured using cosine similarity.

We evaluated the performance of five trajectory inference methods (PAGA [65], Monocle [49], Slingshot [57], MST [51], pCreode [25]) on tree-structured trajectories using the `MT` datasets. The result is shown in Fig. 4b, where we calculated the *R*^2^ and *k*NN purity (Methods) for each separate lineage in each dataset. Overall, PAGA Tree and Slingshot have the best performance, which is in line with results shown in previous benchmarking efforts [51; 69]. When comparing results on datasets with *σ*_cif_ = 0.1 and *σ*_cif_ = 0.5 (Fig. S4a-b), we again see that smaller *σ*_cif_ corresponds to better results. Furthermore, we tested on a simpler linear trajectory dataset `ML1a` (Fig. S4d), and the result was in line with a previous result shown in scDesign3 [54], which used a similar linear trajectory.

### Benchmarking multi-modal and multi-batch data integration methods

A number of computational methods have been proposed to integrate single cell genomics data from multiple modalities and multiple batches [1]. We benchmarked three recently proposed multi-modal integration methods: Seurat bridge integration (Seurat-bridge) [24], UINMF [34] and Cobolt [21] that can integrate data matrices from multiple batches and modalities. We use all 144 main datasets to test their performance under various types of cell population. Each main dataset is divided into three batches (with batch effect 3), then the scRNA-seq data from batch 2 and scATAC-seq data from batch 3 are dropped intentionally to mimic a real scenario where some modalities are missing in certain batches (Fig. 4d). We use the following metrics to evaluate the performance of the integration methods: Adjusted Rand Index (ARI) and Normalized Mutual Information (NMI) as the metrics for cluster identity preservation, and Graph Connectivity and Average Silhouette Width (ASW) as metrics for batch mixing (Methods). These metrics were used in a recent paper on benchmarking single cell data integration methods [40].

The result is shown in Fig. 4c. Since Seurat-bridge does not output the latent embedding for the “bridge” dataset (batch 1 in Fig 4d), only the two matrices from batches 2 and 3 (colored in Fig. 4d) were used for evaluation. We observe that UINMF has the best performance in terms of all four measurements. Seurat-bridge and Cobolt have comparable ARI and NMI but Cobolt has better batch mixing scores. When comparing the ARI and NMI scores for *σ*_cif_ = 0.1 and *σ*_cif_ = 0.5, one can observe that these cell identity preservation scores are higher with smaller *σ*_cif_. Comparing different cell population structures, we see that continuous populations (“Linear” and “Tree”) have lower ARI and NMI scores than discrete populations, potentially because that metrics like ARI and NMI are better suited for discrete populations.

We then ran the integration methods on a large dataset with 3000 cells and visualized the integrated latent embedding in Fig. S5, which helped us to understand each method’s behavior. We noticed that while Seurat-bridge has lower graph connectivity and ASW scores, different batches are located closely (but do not overlap) in the visualized latent space. That the reference and query data in the latent space do not overlap can cause the low batch mixing scores, but may not affect the ability of label transfer.

### Benchmarking RNA velocity estimation methods

We demonstrate scMultiSim’s ability of benchmarking RNA velocity estimation methods by running two representative RNA velocity inference methods, scVelo [3] and VeloCyto [35], on the simulated data. We compare all three models in scVelo package, including the deterministic, stochastic, and dynamical models. The auxiliary dataset `V` (Table 2) was used, which contains 72 datasets of different numbers of cells and genes, with or without GRN. We evaluate the accuracy of inferred RNA velocity using cosine similarity score. The score measures the degree of mismatch between the direction of inferred and ground truth velocity, where a higher score shows a better inference result (Methods).

From the result shown in Fig. 4f, scVelo’s deterministic model has the highest cosine similarity score on all datasets. On the other hand, the dynamical model of scVelo, being considered a generalized version of VeloCyto, does not produce the best result. Interestingly, Gorin et al. also discussed a similar performance issue of the dynamical scVelo. They mentioned that the mismatch between the implicit assumption of dynamical scVelo and the true biological dynamics could be the cause of the performance issue [22]. In spite of the performance differences, the similarity scores are shown to be only around 0.2 for all methods. We suspect that it is the intrinsic noise within the simulated dataset that affect the inference accuracy of all methods. We further conduct experiments comparing the accuracy of inferred RNA velocity with and without *k*NN smoothing (Methods). By using *k*NN smoothing, the inferred RNA velocity of each cell is further averaged with the velocity of all its neighboring cells. Since *k*NN smoothing helps to reduce the noise effect on the inferred velocity, we expect that the overall performance should improve after the smoothing. The experiment results validate our assumption (Fig. 4e), where the average performance of all methods increases to 0.63. The experiments show that the intrinsic noise within the sequencing dataset heavily affects the accuracy of RNA velocity inference methods, and it is still a challenging task to infer RNA velocity from noisy scRNA-seq datasets.

### Benchmarking GRN inference methods

Using scMultiSim, we benchmarked 11 GRN inference methods which were compared in a previous benchmarking paper [48]. Using the predicted networks, we calculate the AUROC (area under receiver operating characteristic curve) as well as the AUPRC (area under precision-recall curve) ratio, which is the AUPRC divided by the baseline value (a random predictor). These metrics were also used in previous benchmarking work [48].

We show results on the 144 main datasets in Fig. 5a. To further inspect the performance in a less-noisy scenario, we also generated auxiliary datasets `G` (Table 2) with a linear trajectory and without CCI effect. We benchmarked the methods using true counts and observed counts in `G`, respectively. The result of `G` is shown in Fig. 5b. All datasets use the same 100-gene GRN from [15]. We observed that PIDC [9] has the best overall performance, especially on true counts. Other methods like GENIE3 [27] and GRNBOOST2 [42] also have noteworthy precision. We then examined the effect of technical noise on the performance of GRN inference methods. On observed counts, both the AUPRC ratio and AUROC value suffer from a decline, indicating that it is significantly harder to infer the GRN from noisy data. However, PIDC continues to have the highest AUPRC and AUROC values, showing that its performance is more resistant to technical noises. SINCERITIES [45], PPCOR [32] and SINGE [14] perform well and beat GENIE3 and GRNBOOST2 on observed counts. Nevertheless, the absolute AUPRC values of all methods, even on true counts, are still far from satisfying, indicating that GRN inference is still a challenging problem.

**Figure 5.**
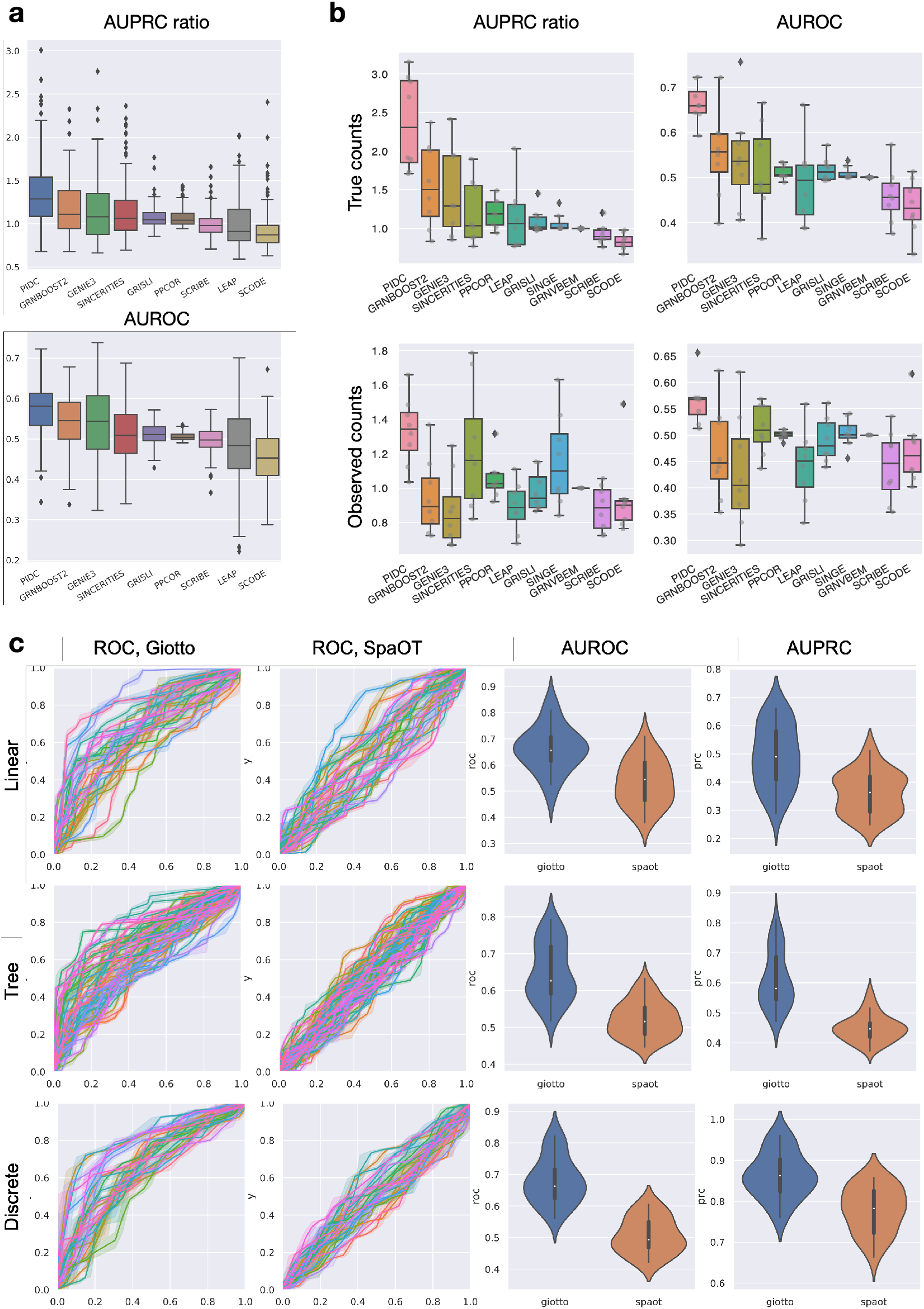
Benchmarking GRN inference and CCI inference methods. (**a**) Benchmarking GRN inference methods. the upper box plot shows AUPRC ratios (versus a random classifier), and the lower box plot shows AUROC values. Methods that did not finish in a reasonable time are excluded. (**b**) Additional results on benchmarking GRN inference methods using datasets G that does not contain CCI effects. Individual data points are also plotted. We also tested the performance on observed counts with technical noise. (**c**) Benchmarking cell-cell interaction inference methods on main datasets with different trajectory types. Each curve in the ROC/PRC plots correspondings to one dataset.

Notably, the ordering of the methods tested using true counts is generally consistent with the ordering reported in [48] even though a different ground truth GRN was used. This fact not only validates the previous results but also suggests that scMultiSim can generate GRN-incorporated gene expression data comparable to other simulators. It indicates scMultiSim’s practicality in benchmarking computational methods that involve GRNs.

### Benchmarking CCI inference methods

Spatially resolved single cell gene expression data provides a powerful tool for understanding cellular processes, tissue organization, and disease mechanisms at the single cell level. Multiple methods have been proposed recently to infer CCIs based on spatial cell locations. However, these methods have yet to be compared in this relatively new field due to the scarcity of biological ground truth and spatial transcriptomics simulators.

We benchmarked three CCI inference methods based on spatially resolved single cell gene expression data, namely Giotto [16], SpaOTsc [5] and SpaTalk [53]. We run Giotto and SpaOTsc on the main datasets and show the result in Fig. 5c. Since SpaTalk needs a minimum of 3 genes from the receptor to a downstream activated TF, we also generated an auxiliary dataset `C` (Table 2) using an artificial GRN with long pathways to satisfy such requirement (Fig. S6a). There are totally eight `C` datasets with 500 cells, 200 genes and a linear trajectory, and the result is shown in fig. S6b. Again, we used AUPRC and AUROC as the metrics. When calculating the PRC and ROC curves, we applied different thresholds on Giotto’s significance score and SpaTalk’s Bonferroni corrected values. Considering both AUROC and AUPRC, Giotto has the best performance with an average AUROC of 0.68 and AUPRC of 0.54 on the main datasets. SpaTalk outputs too many identical p-values for different datasets on dataset `C`, causing the ROC and PRC curves to look unusual. Nevertheless, it has noteworthy performance in terms of AUROC and AUPRC values but is less accurate and stable than Giotto. The benchmarking results show that Giotto could be a versatile yet robust choice for CCI inference.

## Discussion

We presented scMultiSim, a simulator of single cell multi-omics data which is able to incorporate biological factors including cell population, chromatin accessibility, RNA velocity, GRN and spatial CCIs to the output data. We demonstrated the presence of these simulated factors in the generated data, verified the relationship across modalities, and showcased the versatility of scMultiSim through benchmarking on various computational problems. Furthermore, by obtaining consistent benchmarking results with previous works like BEELINE [48] and dyngen [7], the simulated biological effects are validated to be practical and ready for real-world use.

Compared to existing simulators that mainly model one or two biological factors, scMultiSim generates data with more biological complexity similar to real data. This additional complexity enables researchers to better estimate the real-world performance of their methods on noisy experimental data. Furthermore, with the coupled data modalities in the output, researchers can benchmark computational methods that use multiple modalities, which was previously impossible.

scMultiSim’s extensibility and versatility are central to its design, making it easy to include more biological factors and modalities in its simulations. For example, the framework used to model chromatin regions (RIV) and genes (GIV) can also be extended to include other data modalities, such as the protein abundance data. Additionally, we have shown that our CIF/GIV model is versatile enough to mathematically represent the effects of various biological mechanisms like GRNs and CCIs. In addition to the standard functions of scMultiSim, the model can be expanded to consider more realistic scenarios. For instance, the GRN can be set to a cell-specific and cell-type-specific mode, allowing for a more precise simulation of regulatory interactions. Moreover, the scATAC-seq data and scRNA-seq data can follow different trajectories or clustering structures, while the cell clusters can form less regular shapes than the current convex shapes.

scMultiSim’s usability is supported by several key features. First, it requires minimal and easy-to-construct input. For example, users do not need to prepare a backbone for the trajectory to control the cell population; instead, only a plain R phylogenetic tree or a text file with the Newick format tree is needed. Second, scMultiSim has transparent parameters that are self-explanatory and have a clear effect on the result. The user explicitly sets crucial metrics such as the number of cells and genes. Third, scMultiSim’s separated biological effects provide great flexibility. For example, the GRN can affect cell population shapes, but obtaining the desired trajectory using GRN alone is difficult without explicit control of the cell population. scMultiSim’s diff and non-diff CIF mechanism allows users to set the trajectory to any shape without affecting the GRN effects. Users can also let the GRN control the trajectory by increasing the number of non-diff CIF.

We underline that scMultiSim’s major advantage is its ability to encode various factors into a single versatile model, thus creating a comprehensive multi-modal simulator that can benchmark an unprecedented range of computational methods. More importantly, the coupled data modalities in the output jointly provide more information than a single modality alone, making it ideal for designing and benchmarking new methods on multi-omics data. We believe that scMultiSim has the potential to be a powerful tool for fostering the development of new computational methods for single-cell multi-omics data. Moreover, as more benchmarks are conducted, it can help researchers select the appropriate tool based on the type of data they are working with, leading to more accurate and reliable analyses.

## Methods

### A. The Beta-Poisson model and intrinsic noise

The master equation of the kinetic model represents the steady state distribution of a gene’s expression level given its kinetic parameters, *k*_*on*_, *k*_*off*_, and *s* [43]. The Beta-Poisson model was shown to be equivalent to the master equation [31] with faster calculation. The gene expression level *x* (which is also the mRNA count) can be sampled from the following distribution:

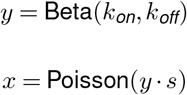

Using the above Beta-Poisson distribution to generate the gene expression level is one mode to obtain mRNA count for a gene in a cell. This works if we only need to generate the spliced mRNA counts. If users also need to generate unspliced mRNA counts and RNA velocity, the other mode, called the “full kinetic model” is used. The Beta-Poisson model is used by default when only generating spliced counts for lower running time.

The sampling process from the Beta-Poisson distribution to obtain *x* introduces intrinsic noise to the data, which corresponds to the intrinsic noise in real data caused by transcription burst. The theoretical mean of the kinetic model, which is 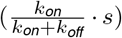, corresponds to the gene expression level of the gene with no intrinsic noise. We introduced parameter *σ*_*i*_ which controls the intrinsic noise by adjusting the weight between the random samples from the Poisson distribution and the theoretical mean:

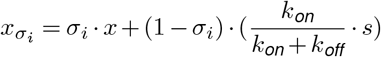

The intrinsic noise in scRNA-seq data is hard to reduce in experiments due to the snapshot nature of scRNA-seq data. The parameter *σ*_*i*_ allows users to investigate the effect of intrinsic noise on the performance of the computational methods.

### B. Cell Identity Factors (CIFs) and Gene Identity Vectors (GIVs)

The length of the CIF and GIV, denoted by *n*_cif_, can be adjusted by the user. Overall, we have a *n*_cell_ *× n*_cif_ CIF matrix for each kinetic parameter (Fig. S1a), where each row is the CIF vector of a cell. Correspondingly, we also have the *n*_cif_ *× n*_gene_ Gene Identity Vectors (GIV) matrix, (Fig. S1b) where each column is linked to a gene, acting as the weight of the corresponding row in the CIF matrix, i.e. how strong the corresponding CIF can affect the gene. In short, CIF encodes the *cell identity*, while GIV encodes the *strength of biological effects*. Therefore, by multiplying the CIF and GIV matrix, we are able to get a *n*_cell_ *×n*_gene_ matrix, which is the desired kinetic parameter matrix with the cell and gene effects encoded. Each cell has three CIF vectors corresponding to the three kinetic parameters *k*_*on*_, *k*_*off*_, and *s*, and similarly for the GIV vectors (Fig. S1a-b).

### C. diff-CIF generates user-controlled trajectories or clusters

When generating data for cells from more than one cell type, the minimal user input of scMultiSim is the cell differentiation tree, which controls the cell types (for discrete population) or trajectories (for continuous population) in the output. The generated scRNA-seq and scATAC-seq data reflect the tree structure through the diff-CIF vectors. The diff-CIF vectors are generated as follows: starting from the root of the tree, a Gaussian random walk along the tree (Fig. 2a) is performed for each cell to generate the *n*_diff-CIF_ dimension diff-CIF vector. Parameter *σ*_cif_ controls the standard deviation of the random walk, therefore a larger *σ*_cif_ will produce looser and noisier trajectory structures. Another parameter *r*_*d*_ is used to control the relative number of diff-CIF to non-diff-CIF. With a larger *r*_*d*_, trajectories are clear and crisp in the output; with a smaller *r*_*d*_, the trajectory is vague, and the shape of the cell population is more controlled by other factors like GRN. For a discrete population, only the cell types at the tree tips are used; then cells of each type are shifted by a Gaussian distribution, controlled by the same *σ*_cif_ parameter. Therefore, a smaller *σ*_cif_ will produce clearer cluster boundaries.

For a heterogeneous cell population, cells have different development stages and types. Users should input a cell differentiation tree where each node represents a cell type. The tree provides a backbone for the trajectory in the cell population. Each dimension of the diff-CIF vector is sampled along the tree via browning motion. First, cells start at the root of the tree; then for each dimension, the diff-CIF value for all cells **v** is

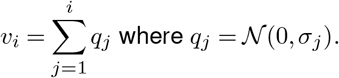

*σ*_*j*_ is the distance along the tree between cell *j* and *j ≠* 1. Alternatively, users can use an impulse model (using the implementation in SymSim). The lengths of the non-diff-CIF and diff-CIF vectors can be controlled by the user. More diff-CIFs will result in a more clear trajectory pattern in the cell population, which corresponds to the input tree. With very few diff-CIFs, the cell population is mainly controlled by the GRN.

### D. tf-CIF and GIV encode the GRN effects

To encode GRN effect in the simulated single cell gene expression data, the GIVs and CIFs are designed to include a “TF part” (Fig. S1a). Cells are generated one by one along the given cell differentiation tree, where the expressions of the TFs in the *t*^th^ cell affect the gene expression of cell *t* + 1. Formally, the *i*^th^ position of the TF part (corresponding to the *i*^th^ TF) of in the CIV of cell *t* +1 is calculated as:

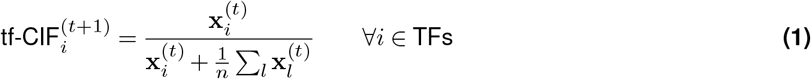

where 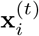 is the expression level of the *i*^th^ TF in the *t*^th^ cell. The corresponding tf-CIF for the root cell is sampled randomly from the Gaussian distribution *𝒩* _cif_ supplied by the user.

The TF part of the GIV for a gene also has length of *n*_TF_(Fig. S1b). Considering all genes, we have a *n*_gene_ *× n*_TF_ matrix, which we call the GRN effect matrix. This matrix encodes the ground truth GRN that is supplied by the user. Naturally, the GRN effect matrix is included in the GIV when calculating the *s* parameter, where the value at (*i, j*) is the regulation strength of TF *j* on gene *i*. Therefore, a larger regulation strength will lead to higher *s*, and consequently, higher expressions for the target genes. For *k*_*on*_ and *k*_*off*_, the tf-CIF vector is sampled using *N*_cif_, assuming that the GRN does not affect the active state of a gene. However, in the scenario where it is desired to model GRN effect also in *k*_*on*_ and *k*_*off*_, similar GRN effect matrix for *s* can be used for *k*_*on*_ and *k*_*off*_.

scMultiSim also allows the use of ground truth GRNs which are cell specific. In this mode, random GRN edges are generated or deleted gradually along the pseudotime at a user-controlled speed. When simulating each cell, the tf-GIV will be filled with the current GRN effect matrix. The cell-specific GRN ground truth is outputted in this mode.

### E. lig-CIF and GIV encode cell-cell interactions

When simulating spatial transcriptomics data with CCI effects, we used a 2-D *k × k* grid to model the spatial locations of cells (Fig. S1d). The grid size *k* is large enough to accommodate the *n* cells (can be specified by the user; if not provided, use 250% of cell number by default). A cell can have at most *n*_nbs_ neighbors with CCI (within the blue circle’s range in Fig. 2a, and this radius can be adjusted). Therefore, the ligand CIF and GIV are of length *n*_lig_ *· n*_nbs_, where *n*_lig_ is the number of ligands.

The lig-GIV vector contains the CCI strength values, for example, the “n2 lg3” entry in Fig. 2a indicates how strong the ligand 3 from the neighbor at position 2 can affect the receptor 2 of this cell. The lig-CIF of each cell will inherit from its previous cell during the simulation process, which is similar to the tf-CIF mentioned above. Each entry of the lig-CIF vector corresponds to a ligand from one neighbor. The same Gaussian distribution *N*_cif_ is used for *k*_*on*_ and *k*_*off*_. For *s*, due to the similarity of the ligand-receptor pairs and the TF-target pairs, we use a similar strategy as tf-CIF (shown in Eq. 1): cell *i*’s lig-CIF is the normalized vector of cell *i ≠* 1’s gene expression counts of the ligand genes (See Fig. 2a, Fig. S1).

At each step *t*, a new cell is considered to be born and added to the grid. When adding a new cell, it has a probability of *p*_*n*_ to be a neighbor of an existing cell with the same cell type. We also provide other strategies to place a new cell, including (1) all cells placed at a random location, and (2) only the first *m* cells are randomly placed, and the remaining follow *p*_*n*_. A pre-defined cell differentiation tree is required as input to define the differentiation topology in the cells. A new cell will always be in the initial state at the root of the differential tree. At each step, an existing cell moves forwards along a random path in the cell differential tree, representing the cell development. The gene expressions in the final step are output as the observed data. Fig. S1 shows the structure for the CCI mode.

To generate ground truth CCIs both at the cell types level and single cell level, scMultiSim pre-defines a ligand-receptor database, represented by a user input *m ×* 3 matrix *S*. There are *m* ligand-target pairs in total that correspond to each row of *S*. For each pair *i*, there are three parameters: ligand gene *L*_*i*_, receptor gene *T*_*i*_, and the effect *E*_*i*_, representing how strongly the ligand can affect the expression of the receptor. For each cell type pair, the ground truth CCI beetween these two cell types are sampled from the ligand-receptor database (corresponds to the columns in *S*). For each neighboring cell pair, the ground truth CCIs between them follow the cell-type-level ground truth CCIs: if the two cells belong to two cell types *C*_1_ and *C*_2_ respectively (where *C*_1_ can be the same as *C*_2_), the CCIs between these two cells follow the CCIs defined in *S* corresponding to pair (*C*_1_, *C*_2_). Users can have further fine-grained control for each cell pair by letting it use a subset of ligand-receptor pairs sampled from the cell-type level ground truth.

Although we collect cells at the last time point as our output (which is the case for real data), different cell types are guaranteed to present in the last step since the cells are added at different time steps, therefore having different development stages. In addition, we let the same cell (at the same location) have the same diff-CIF across different time steps, so the trajectory encoded in the diff-CIF is preserved in the final step. A cell’s TF and ligand CIF for the current step is inherited from the previous one to make sure other factors stay the same.

We use the following steps to calculate the correlation between the expressions of neighboring cells in Fig. 3e. First, a specific ligand-receptor pair (*l, r*) is chosen. Let *T* (*a, b*)= {true, false} denote that there is CCI between cell *a* and cell *b* for (*l, r*). Then, for each cell *i*, we get its neighbor list *n*_*i*_, which is a vector of 4 cells. A vector of 4 non-adjacent cells *m*_*i*_ is also randomly sampled for this cell. Thus, let 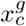 denote the gene expression of cell *c* and gene *g*. we calculate the “neighbor cells with CCI” correlation using the pairs {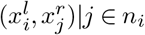,*T* (*i, j*)= true}, the “neighbor cells without CCI” correlation using the pairs {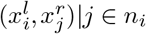,*T* (*i, j*)= false}, and the “non-neighbor cells” correlation using the pairs 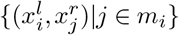. Cell pairs of the same type are ignored while calculating the correlations because they tend to have similar expressions.

### F. Generating the Gene Identity Vectors

A gene’s GIV vector has the same length as the CIF vectors. The values in the GIV of a gene act as the weights of the corresponding factors in the CIF, *i*.*e*., how strong the corresponding CIF can affect the gene (Fig. 2a). If we have *n*_gene_ genes, we obtain a GIV matrix of size *n*_cif_ *× n*_gene_.

It can be divided into four submatrices as shown in Fig. S1b. For *k*_*on*_ and *k*_*off*_, the nd-CIF and diff-CIF are sampled from distribution *𝒢* as shown below:

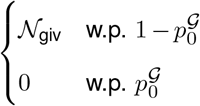

where 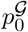 is a parameter specifying the probability of being zero, and 𝒩_giv_ is a user-adjustible gaussian distribution. tf-GIV and lig-GIV are all zeros since TF/ligands affect *s* only. For *s*, the tf-GIV submatrix is the GRN effect matrix, i.e. a *n*_TF_ × *n*_gene_ matrix where the entry at (*i, j*) is the regulation effect between TF *i* and gene *j*. Similarly, the lig-GIV submatrix is the cell-cell interaction effect matrix. The nd-GIV submatrix is sampled from *G*. For diff-GIV, we do the following steps to incorporate the connection between TFs and regulated genes: (1) Randomly select 2 GIV entries for each TF gene and give them a fixed small number. (2) For every target gene, it should use the same GIV vector as its regulators. If a gene has multiple regulators, its gene effects will be the combination of that of the regulators. This is achieved by multiplying the *n*_diff_ *× n*_TF_ GIV matrix in (1) and the *n*_TF_ *× n*_gene_ effect matrix. If a gene is both a TF and target, its GIV will be 0.5 *·* ((1) + (2)).

### G. Simulating scATAC-seq data and relationship between scATAC-seq and scRNA-seq

Since scMultiSim incorporates the effect of chromatin accessibility in the gene expressions, the scATAC-seq data is simulated before the scRNA-seq data. The cell types in the scATAC-seq data can follow the same differentiation tree as in the scRNA-seq data (the scATAC-seq and scRNA-seq data share the same cells) or can follow a different tree (to reflect the difference between modalities).

Similar to GIV, we use a randomly sampled *Region Identity Vector (RIV)* matrix to represent the chromatin regions. Following the same mechanism, we multiply the CIF and RIV matrix, and obtained a “non-realistic scATAC-seq” data matrix. Next, the scATAC-seq data matrix is obtained by scaling the “non-realistic” scATAC-seq data to match a real distribution learned from real data. This is a step to capture the intrinsic variation of the chromatin accessibility pattern, which we will also apply to the kinetic parameters when generating gene expressions.

The RIV matrix is sampled from a distribution ℛ similar to *𝒢*:

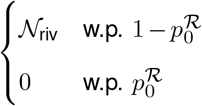

where 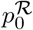 is the probability of being zero and 𝒩_riv_ is a user-adjustable Gaussian distribution. With the CIF and RIV matrices, the *n*_cell_ × *n*_region_ scATAC-seq can be generated by (1) multiplying the CIF matrix by the RIV matrix, (2) scale the matrix to match the real data distribution, and (3) adding intrinsic noise (sampled from a small Gaussian) to the scATAC-seq data. In Step (2), we use the same rank-based scaling process as used for the kinetic parameters as described in Section “Preparing the kinetic parameters” above, and the real scATAC-seq data distribution is obtained from the dataset in [11].

To incorporate the relationship between scATAC-seq and scRNA-seq data, we use the scATAC-seq data to adjust the *k*_*on*_ parameter that is used to generate the scRNA-seq data, considering that chromatin accessibility affects the activated status of genes. First, a region-to-gene matrix (Fig.1b) is generated to represent the mapping between chromatin regions and genes, where a gene can be regulated by 1-3 consecutive regions. Users can input a region distribution vector **r**, for example, (0.1, 0.5, 0.4) means a gene can be regulated by three regions, and the probability of it being regulated by one, two and three consecutive regions are 0.1, 0.5 and 0.4, respectively. The scATAC-seq data is also used to adjust *k*_*on*_ as described in the following section.

### H. Preparing the kinetic parameters

The kinetic parameters, *k*_*on*_, *k*_*off*_ and *s* are needed when generating single cell gene expression data (mRNA counts) using the kinetic model or Beta-Poisson distribution (Fig. 1b). While the basic idea is to get the parameter matrix using CIFs and GIVs (Fig. 1b), the three parameters go through different post-processing after the step of CIF *×* GIV. We first denote the result of CIF *×* GIV for *k*_*on*_, *k*_*off*_ and *s* as *M*_1_, *M*_2_ and *M*_3_, respectively.

i. *k*_*on*_. Since chromatin accessibility controls the activation of the genes, the scATAC-seq data is expected to affect the *k*_*on*_ parameter. We first prepare a *n*_region_ *× n*_gene_ 0-1 region-to-gene matrix *Z* using **r**, where *Z*_*ij*_ indicates region *i* is associated with gene *j* (*Z* is outputted as the region-to-gene matrix). We multiply the scATAC-seq matrix with *Z* to get the *n*_cell_ × *n*_gene_ parameter matrix 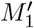. Since the scATAC-seq data is sparse, there are many zeros in 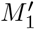. Thus, we replace the zero entries in 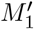 with the corresponding entries in *M*_1_ (scaled to be smaller than the smallest non-zero entry in 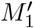) to help differentiate the zero entries. Finally, 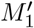 is sampled to match the distribution of *k*_*on*_ inferred from real data.
ii. *k*_*off*_. The parameters are obtained by scaling *M*_2_ to match the real data distribution. For both *k*_*on*_ and *k*_*off*_, it is possible to adjust the bimodality of gene expressions [69] through an optional bimodal factor *B*. A larger *B* will downscale both *k*_*on*_ and *k*_*off*_, therefore increasing the bimodality.
iii. *s*. The parameters are obtained by scaling *M*_3_ to match the distribution of *s* inferred from real data. Then, users can also use a “scale.s” parameter to linearly scale *s*. It allows us to adjust the size of cells – some datasets may tend to large cells and some tend to have small cells depending on the cell types being profiled.

When scaling a matrix (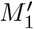, *M*_2_, or *M*_3_) to match a reference distribution (eg. the distributions of *k*_*on*_, *k*_*off*_ and *s* estimated from real data), the procedure is as follows: denoting the reference distribution by *𝒟*, the matrix to rescale by *X*, and the number of elements in *X* by *n*, we sample *n* ordered values from *𝒟*, then replace the data in *X* using the same order. scMultiSim uses the reference kinetic distribution parameters provided in SymSim [69], where the kinetic parameters are estimated from real data via an MCMC approach. The data used are the UMI-based dataset of 3005 cortex cells by Zeisel et al. [67], and a non-UMI-based dataset of 130 IL17-expressing T helper cells (Th17) by Gaublomme et al [20].

### I. Generating RNA velocity with the full kinetic model

When using the full kinetic model, scMultiSim can generate the spliced and unspliced counts for each cell from the kinetic parameters. The starting spliced count *x*_*s*_ and unspliced count *x*_*u*_ for a cell are the previous cell’s counts on the differential tree. For the first cell, the spliced/unspliced counts are

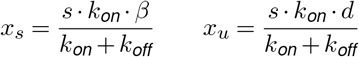

where *β* and *d* respectively represent the splicing and degradation rate of genes. Both *γ* and *d* are sampled from a user-controlled normal distribution.

We set the cell cycle length to be 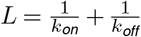, and divide it into multiple steps. The number of steps follows 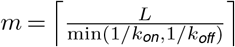. We also provide an optional cell length factor *÷*_*L*_ parameter to scale the cycle length. The probabilities of gene switching on or off are then calculated with 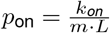 and 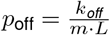. In each simulation step, we update the cell’s current on/off state based on *p*_on_ and *p*_off_, and generate the spliced/unspliced counts *x*_*s*_ and *x*_*u*_. The spliced counts at step *t* are obtained by:

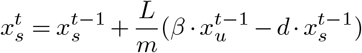

and the unspliced counts are obtained by:

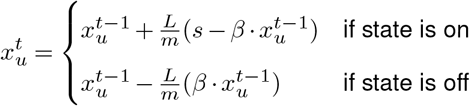

The outputted *x*_*s*_ and *x*_*u*_ are the values at the final step *t* = *m*. The ground truth RNA velocity is calculated as:

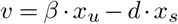

We obtain the KNN averaged RNA velocity by applying a Gaussian Kernel KNN on the raw velocity data, with *k* = ⌈*n*_cell_*/*50⌉. Then we normalize the velocity by calculating each cell’s normalization factor *s*_*i*_ = |*v*_*i*_|, where *v*_*i*_ is the velocity vector for cell *i*.

### J. Adding technical noise and batch effects to data

Technical noise is added to the true mRNA counts to generate observed counts (observed scRNA-seq data) (Fig. 1b). The workflow follows SymSim’s approach [69]: we simulate multiple rounds of mRNA capture and PCR amplification, then sequencing and profiling with UMI or non-UMI protocols. The parameter *–* controls the capture efficiency, that is, the rate of subsampling of transcripts during the capture step, which can vary in different cells, and user can specify it using a Normal distribution *α ∼ 𝒩* (*α*_*µ*_, *α*_*σ*_). The sequencing depth *d ∼ 𝒩* (*d*_*µ*_, *d*_*σ*_) is another parameter that controls the quality of the observed data.

Batch effects are added by first dividing the cells into batches, then adding gene-specific and batch-specific Gaussian noise based on shift factors. For each gene *j* in batch *i*, the shift factor is sampled from Unif(*µ*_*j*_ *− e*_*b*_, *µ*_*j*_ + *e*_*b*_), where *µ*_*j*_ *∼ 𝒩* (0, 1), and *e*_*b*_ is the parameter controlling the strength of batch effects. We provide several settings for adding highly expressed genes to help researchers fit the housekeeping genes in real data. scMultiSim also supports cell- and gene-wise tuning of the mRNA capture efficiency during the PCR process; therefore per-cell and per-gene metrics (such as zero count proportion and count variance) in the observed data can be controlled separately.

For scATAC-seq data, as the data is sampled from real data we do not explicitly simulate the experimental steps. We do provide methods to add batch effects to obtain multiple batches of scATAC-seq data.

### K. Comparing statistical properties of simulated data with experimental data

To measure scMultiSim’s ability to generate realistic data while incorporating all the effects, we compare the statistical properties of a real mouse somatosensory cortex seqFISH+ [19] dataset with simulated data generated using selected parameters. The dataset, with 10000 genes and spatial locations of 523 cells, is featured in Giotto [16]’s tutorial.

The scMultiSim simulated data has both GRN and CCI effects. The GRN used as input to scMultiSim is obtained as follows: GENIE3 [27] was used to obtain an inferred GRN from the dataset, then after looking at the output edge importance values, the top 200 edges were utilized to form a reference GRN. We used this GRN (96 genes) and another randomly sampled 104 genes to generate a subsample of the data. We then simulated a dataset with 200 genes and 523 cells using scMultiSim. After observing the dimension reduction of the real dataset, a discrete cell population is assumed. We specify the cluster ground truth using the exact cell type labels in the dataset. There are 10 cell types in total. We also used Giotto [16] to infer the cell-cell interactions between cells. We chose the top-seven most significant ligand-receptor pairs from Giotto’s output, with p-value *Æ* 0.01, more than 10 ligand and 10 receptor cells, and the largest `log2fc` values.

We used dyngen [7] as a baseline simulator to compare with scMultiSim. We generated a simulated dataset with dyngen, using the same GRN and number of cells. The cell types and cell-cell interaction ground truth were not provided since dyngen does not support them. Yet, we supplied the raw mouse SS cortex count matrix to dyngen’s `experiment_params` as a reference dataset.

We used the following metrics to compare the distribution of simulated and experimental datasets, which is also used in [15]: library size (per cell), zero counts proportion (per cell), zero counts proportion (per gene), mean counts (per gene), counts variance (per gene), and the relationship between zero counts and mean counts per gene.

### L. Evaluation metrics for benchmarking computational methods

When evaluating the trajectory inference methods, we calculate the coefficient of determination *R*^2^ and the *k*NN purity for all cells on each lineage. Given the cells’ ground truth pseudotime vector *t* and the inferred pseudotime 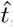, the *R*^2^ is equal to the square of the Pearson correlation coefficient:

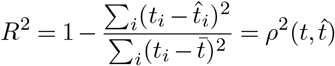

where 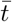 is the mean of *t*. Given a cell *i*’s *k*NN neighborhood 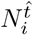 in 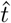 and its *k*NN neighborhood 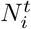 in *t*, the *k*NN purity *K*_*p*_ for the cell is the Jaccard Index of 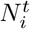 and 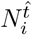.

The evaluation metrics used for multi-model data integration methods, Graph Connectivity and ASW, are described as following.

Graph Connectivity is defined as:

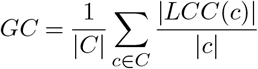

where *C* is all cell types, *LCC*(*c*) is in the largest connected component for cells of type *c*.

For the ASW:

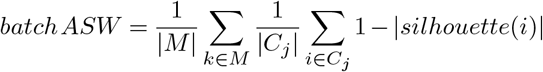

where *M* is the set of all cell types, and *C*_*j*_ is all the cells of type *j*. We used the implementation in [40].

When evaluating RNA velocity inference methods, we used the *cosine similarity* between the averaged estimated velocity and the ground truth. Calculating the average of estimated velocity vectors is commonly used to reduce local noise [7]. In dyngen [7], averaged RNA velocities were calculated across cells at trajectory waypoints weighted through a Gaussian kernel using ground truth trajectory; while in scMultiSim, we averaged the raw velocity values by *k*NN with a Gaussian kernel and *k* = *n*_cells_*/*50 to achieve a similar averaging effect. Finally, cosine similarity is calculated as:

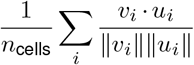

where *v*_*i*_ is the ground truth velocity vector for cell *i*, and *u*_*i*_ is the predicted velocity vector.

### M. Details on running clustering methods

We used CIDR 0.1.5, SC3 1.24.0, Seurat 4.1, and TSCAN 2.0. The parameters we specified are (1) SC3: `pct_dropout` = [0, 100], (2) Seurat: `dims.use` = 30. For PCA-Kmeans, we simply ran Kmeans clustering on the first 20 principle components using the default R implementation `prcomp` and `kmeans`. ARI is calculated by `adjustedRandIndex` from the R package `mclust`. Some code was adapted from [17].

### N. Details on running trajectory inference methods

We used the latest dynverse [51] package (dyno 0.1.2) to run the trajectory inference methods. When running them, we provide the correct root cell ID, number of starting clusters and number of ending clusters. The *R*^2^ values are calculated between the inferred pseudotime and the ground truth for each separate lineage. The *k*NN purity value is calculated for each lineage as: for cell *i*, we obtain its *k* Nearest Neighbors *N*_*i*_ on the pseudotime with *k* = 50. Then the *k*NN purity for *i* is the Jaccard Index of *N*_*i*_ on the inferred pseudotime and *N*_*i*_ on the true pseudotime. *R*^2^ measures the correctness of inferred pseudotime, but when there are multiple branches in the trajectory, *R*^2^ does not distinguish cells with similar pseudotime but are on different branches. In this case, the *k*NN purity serves as a complementary measurement that measures the correctness of inferred trajectory backbone.

### O. Details on running data integration methods

We use all 144 main datasets. Technical noise and batch effects were added using default parameters (non-UMI, *α ∼ 𝒩* (0.1, 0.02), depth *∼ 𝒩* (10^5^, 3000), ATAC observation probability 0.3). All integration methods were run on the scRNA and scATAC data with technical noise and batch effect. For Seurat-bridge, we followed the vignette “Dictionary Learning for cross-modality integration” in Seurat 4.1.0 using the default parameters. For UINMF, we used the latest GitHub release. We followed the “UINMF integration of Dual-omics data” tutorial and ran the `optimizeALS` method using *k* = 12. For Cobolt, we used the GitHub version cd8015b, with 10 latent dimensions, learning rate 0.005. If the loss diverged, we automatically retry with learning rate 0.001. The metrics, including ARI, NMI, Graph Connectivity, and ASW were computed using the scib [40] package.

### P. Details on running RNA velocity estimation methods

We use the datasets `V` to benchmark RNA velocity inference methods as shown in Table 2. We used scVelo 0.2.4 and VeloCyto 0.17.17. We benchmarked scVelo with three modes: `deterministic`, `stochastic`, and `dynamical`. For VeloCyto, we used the default options.

### Q. Details on running GRN inference methods

We use the BEELINE [48] framework to benchmark GRN inference methods. Apart from the main datasets, The dataset `G` (Table 2) was generated using the following configurations: The 100-gene GRN in Fig. 3, 1000 cells, 50 CIFs, *r*_*d*_ = 0.2, *σ*_*i*_ = 1, with other default parameters. Eight datasets were generated for random seed 1 to 8, and technical noise and batch effect was added using default parameters. We ran the BEELINE GitHub version 79775f0. In order to resolve runtime errors, all docker images were built locally, except that we used the provided images on Docker Hub for PIDC and Scribe. We use BEELINE’s example workflow to infer GRN and calculate the AUPRC ratio and AUROC for (a) true counts in the eight datasets, and (b) observed counts with batch effects in the eight datasets. The AUPRC ratio is the AUPRC divided by the AUPRC of a random predictor, which equals to the network density of the ground truth network. Eleven methods were benchmarked in total: PIDC, GRNBoost2, GENIE3, Sincerities, PPCOR, LEAP, GRISLI, SINGE, GRNVBEM, Scribe and SCODE. Methods that did not finish in a reasonable time are excluded in the figure.

### R. Details on running CCI inference methods

We generated 12 datasets using the following procedure. Apart from the main datasets, for each `C` dataset (Table 2), we first construct the GRN (Fig. S6a): (1) let genes 1-6 be the transcription factors. Sample 70 edges from gene 1-6 to gene 7-53. (2) Connect gene 7-53 (regulator) to gene 54-100 (target) consecutively. (3) Connect gene 54-100 to gene 110-156. In this way, we can generate a GRN with reasonable edge density and make sure that there are three downstream genes for each TF, which is required by SpaTalk. Then we construct the ligand-receptor pairs: let the ligands be gene 101-106 and receptors be gene 2, 6, 10, 8, 20, and 30. We divide a linear trajectory into 5 sections, corresponding to 5 cell types. Between each cell type pair (excluding same-type pairs), we sample 3-6 ligand-receptor pairs and enable cell-cell interactions with them for the two cell types. The dataset is then simulated using 160 genes in total, 500 cells, and 50 CIFs. We use the true counts to benchmark the methods.

To run SpaTalk, as suggested by the authors, we used the latest GitHub version and modified the original `plot_lrpair_vln` method to return the p-value from the Wilcoxon rank sum test directly, rather than drawing a figure. Before using the p-values to calculate the precision and recall, we adjusted them using Bonferroni correction:

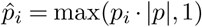

where *p* is the p-value vector for all cell types and ligand-receptor pairs. For Giotto, we used the R package 1.1.2 and followed the `mini_seqfish` vignette. For SpaOTsc, we used version 0.2 with default parameters.

## Data and Code Availability

The scMultiSim R package is available at https://github.com/ZhangLabGT/scMultiSim. The code for dataset generation and benckmarking is available at https://github.com/ZhangLabGT/scMultiSim_manuscript. The simulated datasets are available upon requests.

## Author Contributions

X.Z. conceived the idea and X.C. contributed to the design of scMultiSim. H.L., Z.Z. and M.S. implemented the software package. H.L. performed validations and benchmarks. H.L., X.Z. and Z.Z. wrote the manuscript.

## Acknowledgements

This work was supported by the US National Science Foundation DBI-2019771 and National Institutes of Health grant R35GM143070 (HL, ZZ, XZ), the Guangdong Basic and Applied Basic Research Foundation (2022B1515120077 to XC) and the Shenzhen Innovation Committee of Science and Technology (20220815094330001 to XC).

