## Supplementary Information for "scMultiSim: simulation of multi-modality single cell data guided by cell-cell interactions and gene regulatory networks"

| | | | | |
|---|---|---|---|---|
| **Modalities simulated** | | | | |
| mRNA counts | ✔ | ✔ | ✔ | ✔ |
| Unspliced mRNA counts | ✔ | ✔ | ✔ | |
| Cell spatial location | ✔ | | | ✔ |
| Chromatin accessibility | ✔ | | | ✔ |
| **User-controlled ground truth** | | | | |
| Continuous population - Trajectory structure | ✔ | ✔ *1 | | ○ *2 |
| Continuous population - Trajectory looseness | ✔ | | | |
| Discrete population - Cluster labels | ✔ | | ✔ | ○ *2 |
| Discrete population - Cluster sizes | ✔ | | | ○ *2 |
| Gene regulatory network | ✔ | ✔ *1 | ✔ | *3 |
| Cell-cell interaction network | ✔ | | | |
| **Simulator-provided ground truth** | | | | |
| Cluster labels | ✔ | ✔ | ✔ | ✔ |
| Trajectory | ✔ | ✔ | ✔ | ✔ |
| Cell-specific gene regulatory network | ✔ | | ✔ | |
| Cell-cell interaction (CCI) network | ✔ | | | |
| Chromatin region - gene correspondence | ✔ | | | |
| **Technical Variations** | | | | |
| Single-batch technical noise | ✔ | ✔ | ✔ | ✔ |
| Batch effect | ✔ | ✔ | ✔ | ✔ |
| **Benchmarking computational methods** | | | | |
| Clustering | ✔ | ✔ | ✔ | ○ *4 |
| Trajectory inference | ✔ | ✔ | ✔ | ○ *4 |
| GRN inference | ✔ | ✔ | ✔ | |
| Cell-cell interaction inference | ✔ | | | |
| RNA velocity estimation | ✔ | ✔ | ✔ | |
| Multi-batch data integration | ✔ | ✔ | ✔ | ○ *4 |
| Multimodal data integration | ✔ | | | ○ *5 |
| Mosaic data integration | ✔ | | | |
| Joint-modal network / CCI inference | ✔ | | | |

*1 dyngen assumes the trajectory is determined by the GRN. In order to achieve certain trajectory structures, users need to use specific GRN inputs to obtain the desired trajectory structure of the output data.

*2 In scDesign3, the cluster labels and trajectory structure (pseudotime) used as ground truth are obtained from annotations in the reference data. Users can only modify the ground truth later in the fitted model. The quality of user-provided labels will also affect the similarity between simulated data and its reference data.

*3 scDesign3 does not provide ground truth GRN, but users can modify the learned gene-gene correlations in the fitted model.

*4 Since scDesign3 is reference-based, users need to have an appropriate reference dataset with good-quality annotations. scDesign3 performs model selection based on criteria that do not require the ground truth, e.g., the BIC.

*5 scDesign3 achieves cross-modality coupling by integrating real datasets of different modalities.

**Table S1.** Comparison of scMultiSim and existing multi-modal simulators

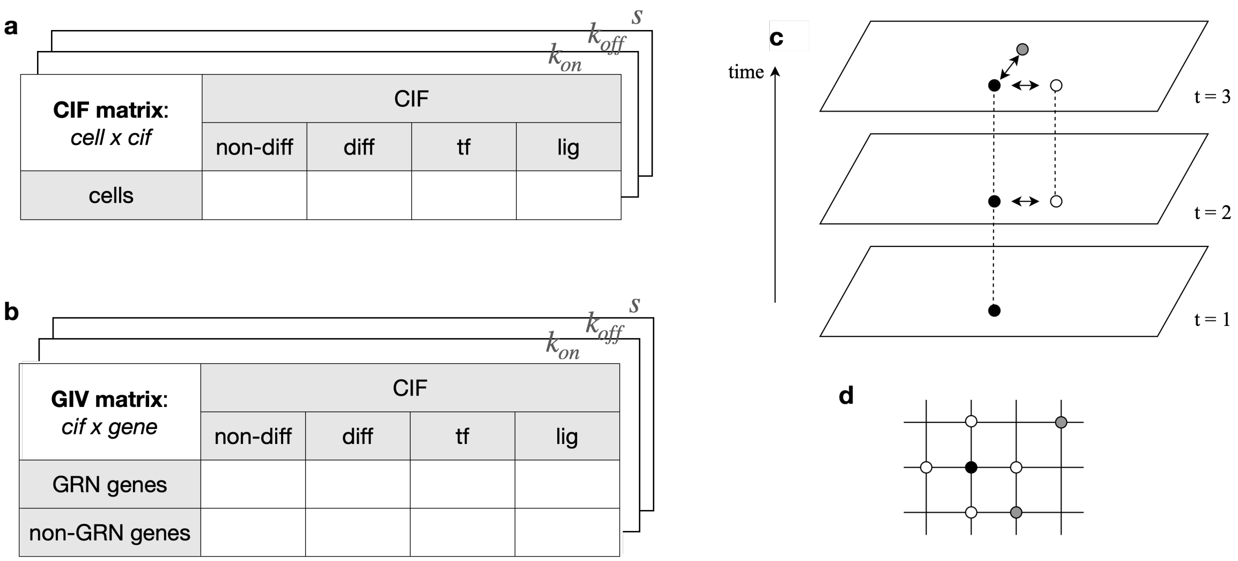


**Figure S1.** **(a)** The CIF matrix of size cell$\times n_{\text{cif}}$. **(b)** The GIV matrix of size $n_{\text{cif}}\times$ gene, transposed for clarity. Its rows match the columns in the CIF matrix, representing the effect (weight) of each gene to those factors. **(c)** We perform the same simulation for $n_{\text{step}}$ steps, adding one new cell in each step. Spatial interactions in each step are incorporated. **(d)** A cell (black) and its neighbors (white) in the grid. The cells in grey are not neighbors.

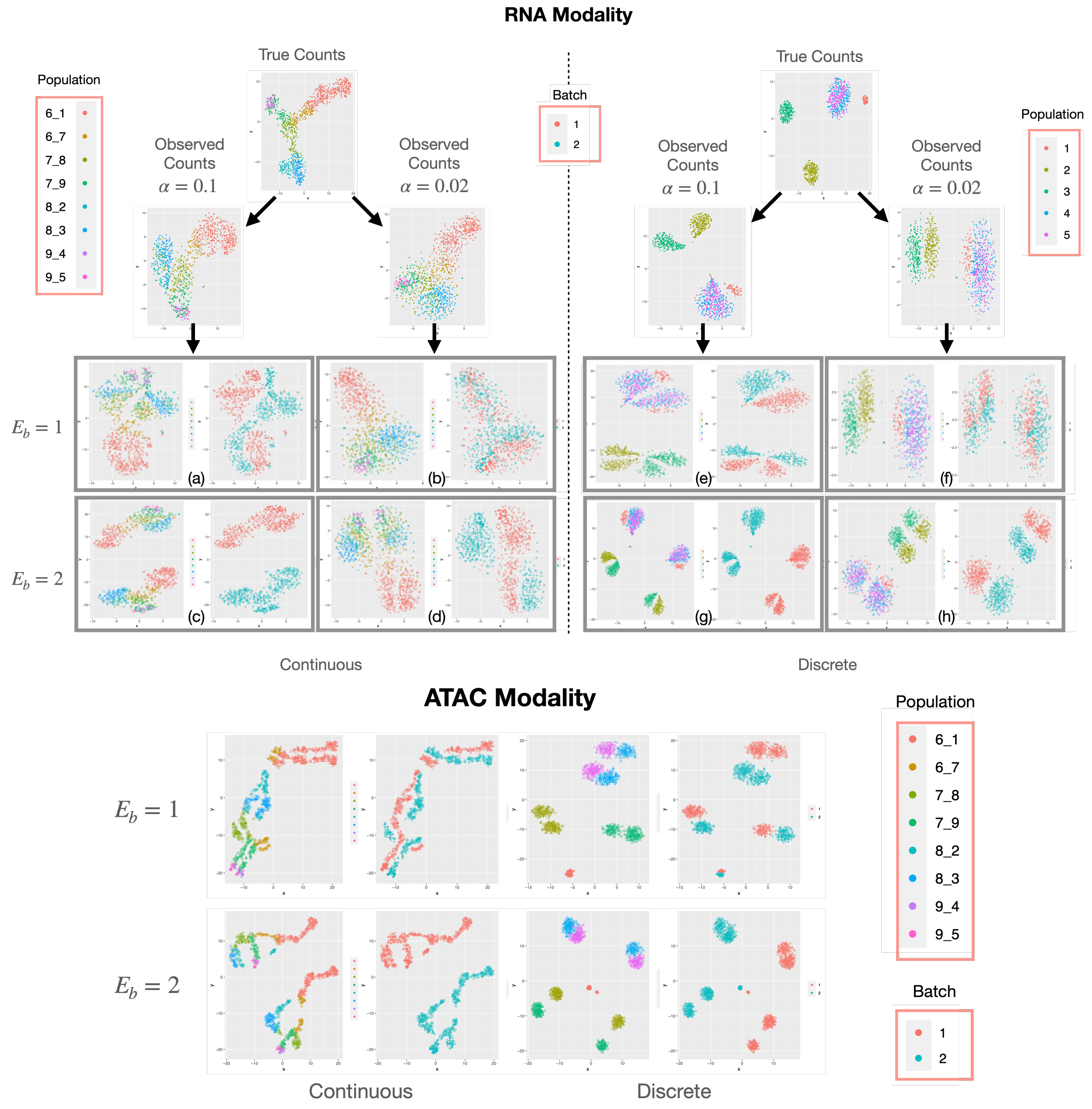


**Figure S2. Additional results on technical variation with different capture efficiency $\alpha$ and batch effect size $E_b$.** Data was simulated using the tree in Fig.2b, with $\sigma_i = 1$, $r_d = 0.2$, 500 genes and 1000 cells. From the same true counts, various technical noise was added for both continuous and discrete cell population. We show the t-SNE visualization of the gene expression under four configurations $\alpha = \{0.1, 0.02\}$, $E_b = \{1, 2\}$, and the chromatin accessibility for $E_b = \{1, 2\}$. In each grey box, the left sub-figure is colored by cell population ground truth, and the right is colored by batches. With a lower capture efficiency $\alpha$, one can easily observe the deterioration of data quality in both discrete or continuous trajectories. For example, cluster 3 in (e) is separated from cluster 4 and 5, while in (f) clusters 3, 4 and 5 cannot be differentiated in the visualization; clusters in (a) also have clearer boundaries than (b). The effect of batch effect $E_b$ is also visible in the visualization, where batches are more separated when $E_b = 2$ in (c,d,g,h). Same observation also applies to the scATAC-seq data.

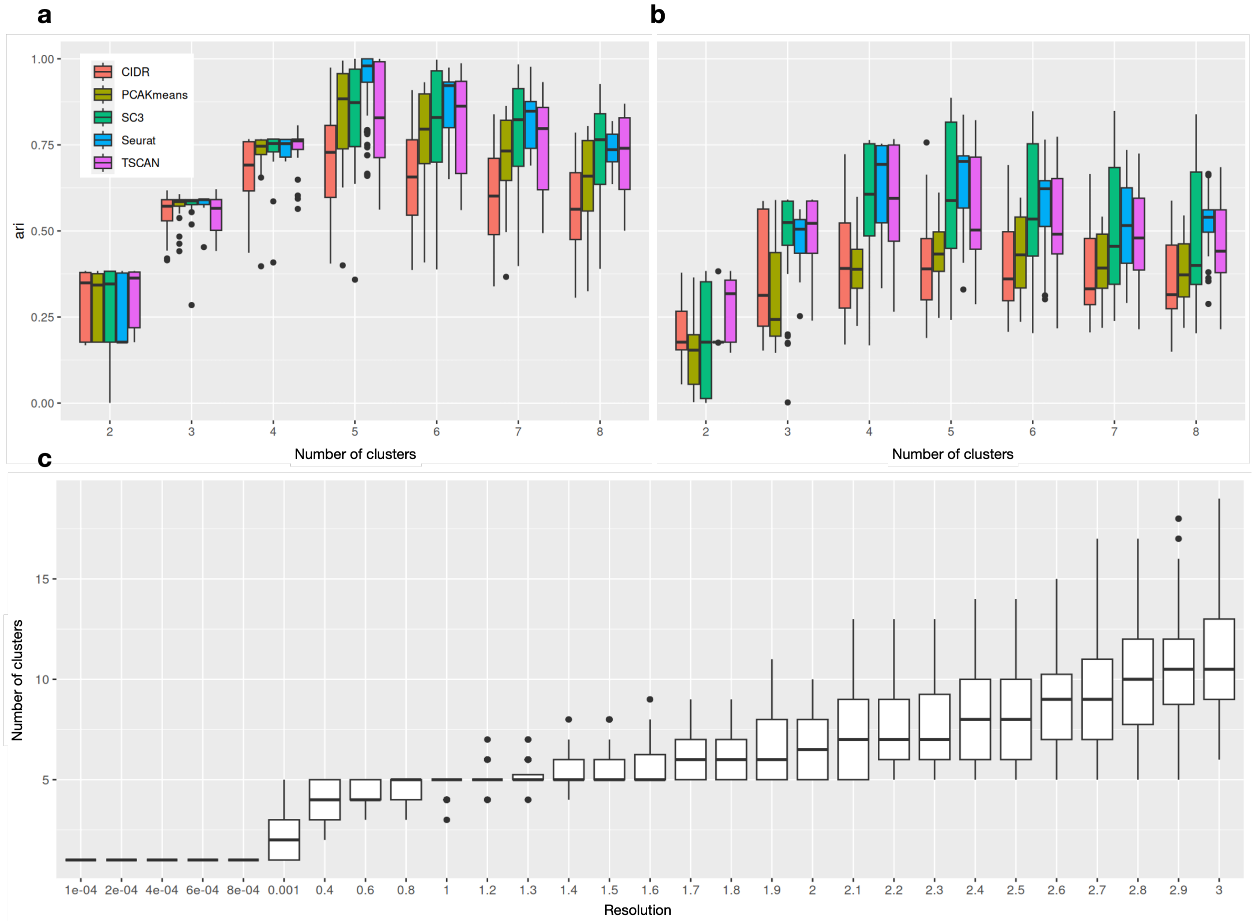


**Figure S3.** Additional results on benchmarking clustering methods. **(a)** Result only using the datasets when $\sigma_{\text{cif}} = 0.1$. **(b)** Result only using the datasets when $\sigma_{\text{cif}} = 0.5$. Performance decrease can be observed when there are more noise. SC3 can perform better in the more noisy cases, while Seurat still provides robust result with good accuracy. **(c)** Relationship between Seurat's resolution parameter and number of clusters in the result. When resolution is between 0.6-1.4, the resulting cluster number is very accurate. This range is in line with Seurat's recommended resolution.

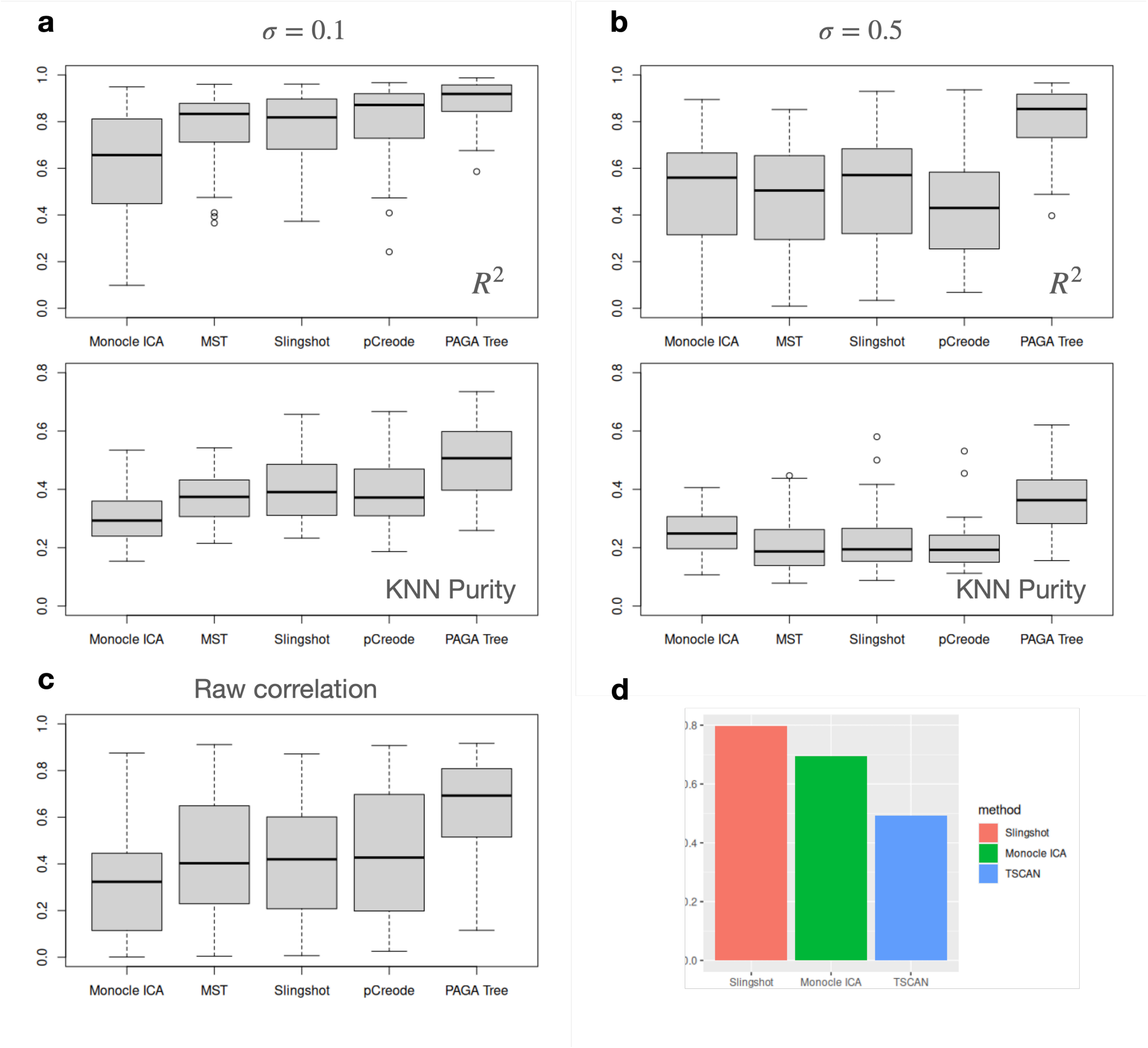


**Figure S4.** Additional results on trajectory inference methods. **(a)** $R^2$ and KNN purity results when $\sigma_{\text{cif}} = 0.1$. **(b)** $R^2$ and KNN purity results when $\sigma_{\text{cif}} = 0.5$. Performance decrease can be observed when there are more noise. **(c)** Pearson correlation calculated directly between the ground truth and the output pseudotime, without averaging on different lineages. **(d)** We also tested the single-lineage curve-fitting performance on dataset `ML1a` , which has a linear trajectory with $\sigma_{\text{cif}} = 0.1$. We tested Slingshot, Monocle ICA, and another popular linear trajectory inference method TSCAN. This result is in line with the result shown in scDesign3 simulated from a dataset originally generated by dyngen.

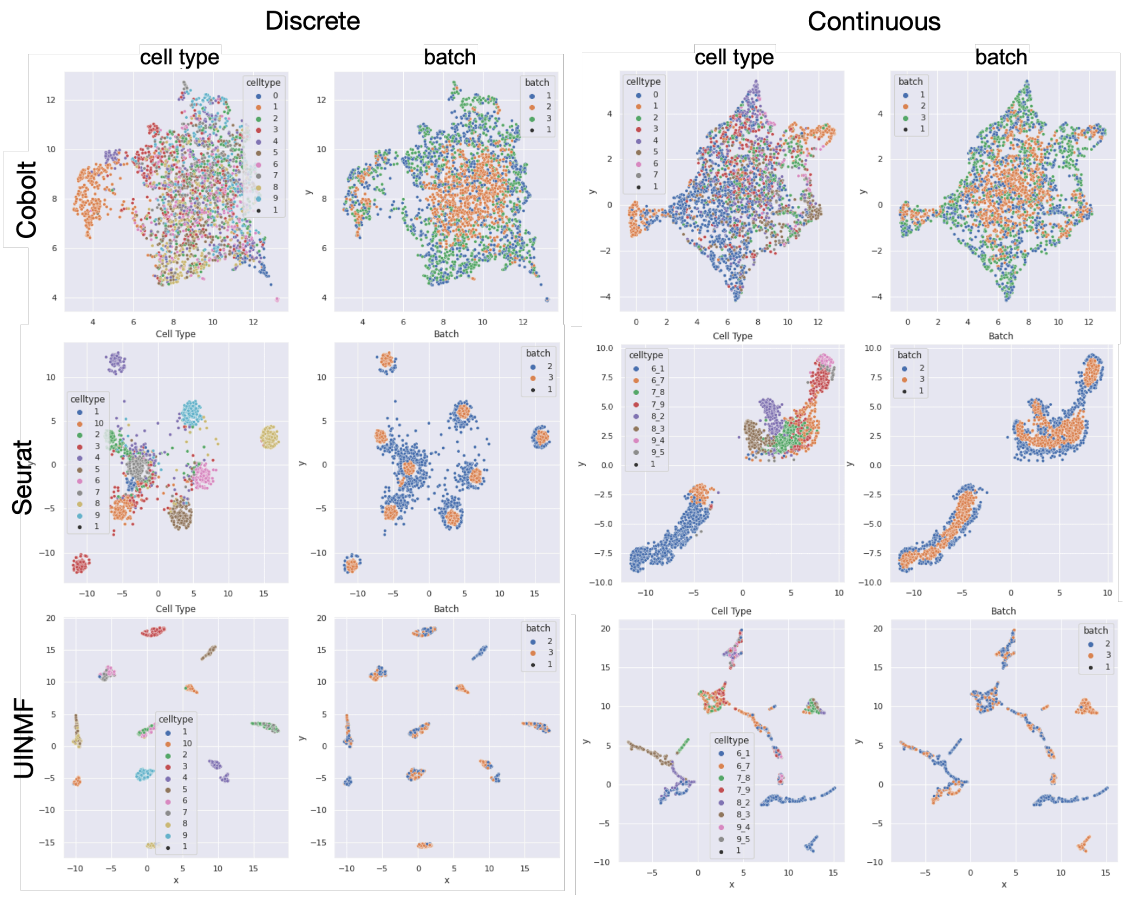


**Figure S5.** Additional results on integration benchmarking. Visualization of each data integration methods's result, using tSNE on a dataset simulated with 3000 cells and 200 genes.

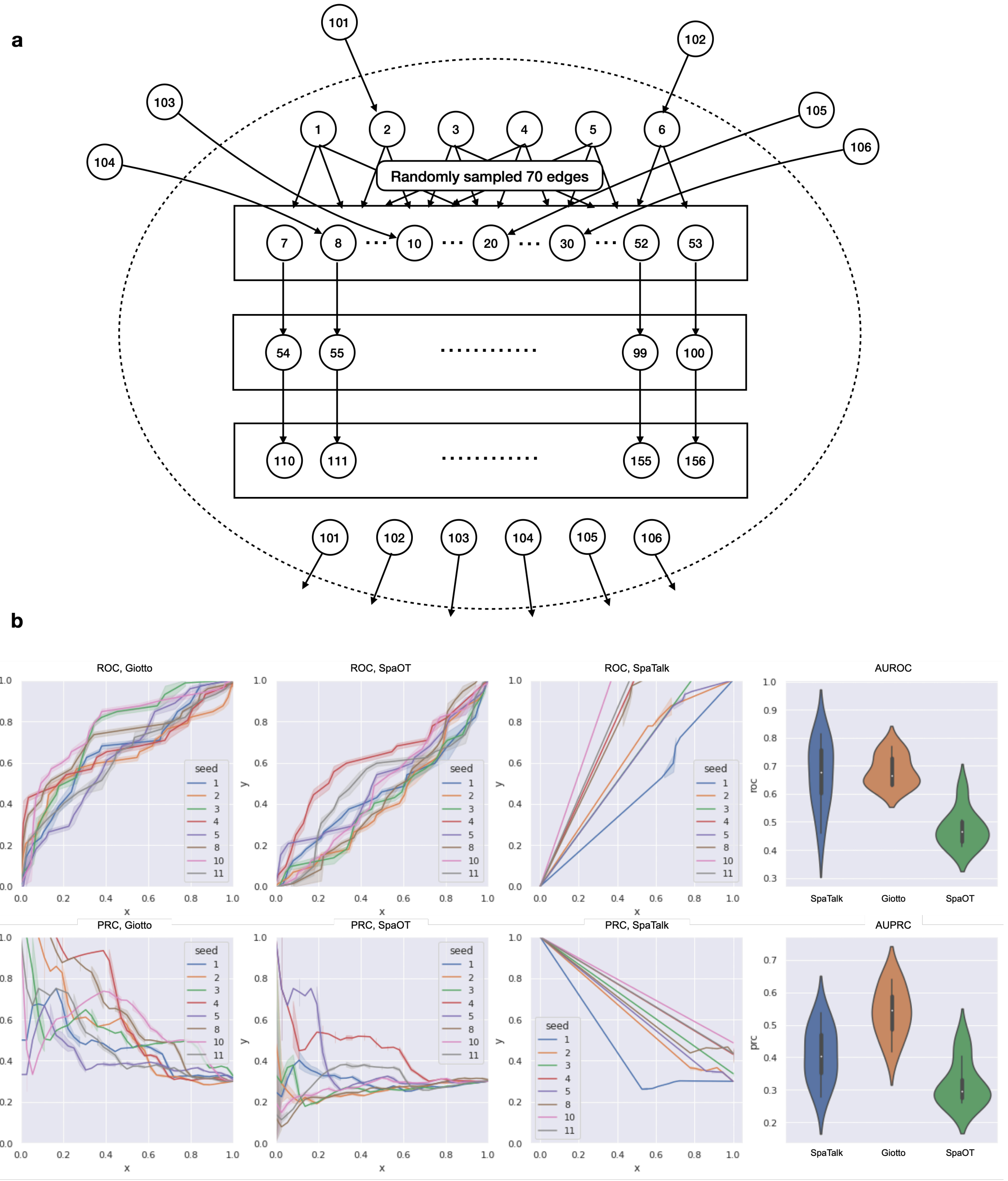


**Figure S6.** Additional results on CCI benchmarking. **(a)** The GRN and CCI network used in datasets C. **(b)** Comparison of Giotto, SpaOTsc, and SpaTalk on dataset C.
